# The protozoan *Trichomonas vaginalis* targets bacteria with laterally-acquired NlpC/P60 peptidoglycan hydrolases

**DOI:** 10.1101/320382

**Authors:** Jully Pinheiro, Jacob Biboy, Waldemar Vollmer, Robert P. Hirt, Jeremy R. Keown, Anastasiia Artuyants, David C. Goldstone, Augusto Simoes-Barbosa

**Author notes:** Corresponding authors: David Goldstone Augusto Simoes-Barbosa.

## Abstract

*Trichomonas vaginalis* is a human eukaryotic pathogen and the causative agent of trichomoniasis, the most prevalent non-viral sexually transmitted infection worldwide. This extracellular protozoan parasite is intimately associated with the human vaginal mucosa and microbiota but key aspects of the complex interactions between the parasite and the vaginal bacteria remain elusive. We report that *T. vaginalis* has acquired, by lateral gene transfer from bacteria, genes encoding peptidoglycan hydrolases of the NlpC/P60 family. Two of the *T. vaginalis* enzymes were active against bacterial peptidoglycan, retaining the active site fold and specificity as DL-endopeptidases. The endogenous NlpC/P60 genes are transcriptionally up regulated in *T. vaginalis* when in the presence of bacteria. The over-expression of an exogenous copy produces a remarkable phenotype where the parasite is capable of competing out bacteria from mixed cultures, consistent with the biochemical activity of the enzyme *in vitro*. Our study highlights the relevance of the interactions of this eukaryotic pathogen with bacteria, a poorly understood aspect on the biology of this important human parasite.

**Author summary:** *Trichomonas vaginalis* is a protozoan parasite that causes a very common sexually transmitted disease known as trichomoniasis. This extracellular parasite resides in the vagina where it is in close association with the mucosa and the local microbiota. Very little is known about the nature of the parasite-bacteria interactions. Here, we report that this parasite had acquired genes from bacteria which retained their original function producing active enzymes capable of degrading peptidoglycan, a polymer that is chemically unique to the cell envelope of bacteria. Our results indicate that these enzymes help the parasite compete out bacteria in mixed cultures. These observations suggest that these enzymes may be critical for the parasite to establish infection in the vagina, a body site that is densely colonised with bacteria. Our study further highlights the importance of understanding the interactions between pathogens and microbiota, as the outcomes of these interactions are increasingly understood to have important implications on health and disease.

## Introduction

*Trichomonas vaginalis* is a flagellated protozoan parasite that causes human trichomoniasis, the most common non-viral sexually transmitted infection worldwide [1]. Women experience symptoms more often than men and these are mostly associated to vaginitis and gyneco-obstetric complications [2–7]. In addition, trichomoniasis facilitates the transmission of the Human Immunodeficiency Virus [8]. *T. vaginalis* is an extracellular mucosal parasite that displays an intimate association with the host tissue and vaginal microbiota. The molecular and cellular basis of the interactions between *T. vaginalis* and this microbiota are poorly understood but likely to have a significant impact on the pathobiology of this infection [9–11]. In addition, trichomoniasis is apparently associated with microbial dysbiosis of the vagina [12].

*T. vaginalis* is thought to have evolved from an enteric to a genitourinary mucosal environment [13] and has been continuously in contact with the mucosa-associated microbiota. Adaptations to colonize the vaginal mucosa may have impacted on (or facilitated by) a recent and dramatic genome expansion achieved through a complex combination of gene and transposable elements duplications and lateral gene transfers (LGTs). These have contributed to the unexpectedly large genome size (~170 Mbp) and the extraordinary coding capacity of ~60,000 predicted protein coding genes [14,15], of which about 30,000 where shown to be transcribed in various tested growth conditions [16,17]. The co-evolution of *Trichomonas* and bacterial members of the microbiota may have significantly shaped the genome of this parasite by providing new functionalities and selective advantages for the parasite to infect human mucosal surfaces of the urogenital tract [15,18,19].

Genome-wide analyses indicate that a considerable number of genes have been acquired by eukaryotes from prokaryotes by LGT [15,20–22]. Among these are an increasing number of genes encoding for enzymes that degrade or remodel the essential bacterial cell wall component, the peptidoglycan (PG) [19,22–26]. Two reports, in particular, have directly demonstrated beneficial implications of these LGT-acquired genes to the recipient eukaryotic organisms [25,26]. The acquisition of bacterial PG-degrading enzymes in these two cases has provided a new function to the eukaryotic host which is controlling bacterial presence or abundance [22,25,26].

In the ecological context of *Trichomonas* and commensal bacteria, lateral acquisition of PG-degrading enzymes could be advantageous to the parasite and it may explain why *T. vaginalis* infections are preferentially accompanied by certain species of vaginal bacteria [12]. Indeed, NlpC/P60-like genes are among strong candidates of LGT among the annotated protein coding genes in the *T. vaginalis* genome [14,18]. NlpC/P60 proteins were originally described as bacterial cell wall endopeptidases cleaving the D-γ-glutamyl-*meso*-diaminopimelate linkage in PG [27,28]. NlpC/P60 proteins display the conserved catalytic triad of papain-like thiol peptidases and may carry additional domains that specify substrate binding, signal peptides or transmembrane regions for proper subcellular localisation [28].

This study aimed to (i) understand the structural diversity and evolutionary origins of NlpC/P60-like genes identified in the *T. vaginalis* genome, (ii) characterize the structure-function relationship for a selection of these enzymes by determining their crystal structures and activities against PG and (iii) examine the potential role of NlpC/P60 enzymes in parasite-bacteria interactions. To the best of our knowledge, this is the first study to report the preservation of fully functional LGT-derived PG-degrading enzymes in a eukaryotic pathogen of the mucosa.

## Results

### Primary structural diversity and phylogeny of NlpC genes from *Trichomonas vaginalis*

A screen for candidate LGT of prokaryotic origins among annotated protein coding genes in the *T. vaginalis* G3 genome identified the entry TVAG_119910 (XP_001276902.1) as a strong candidate LGT from bacterial origin [14,18]. This gene encodes a protein annotated as an endopeptidase member of the Clan CA, family C40, NlpC/P60 (Pfam entry PF00877 - NlpC/P60 family). A total of nine annotated NlpC/P60 proteins [14] with the domain PF00866 were identified in TrichDB [29] (S1 Table and Fig 1A and S1 Fig), referred to as TvNlpC/P60. InterProScan analyses of these proteins also identified a bacterial SH3 domain (domain PF08239, SH3b) in two of the TvNlpC/P60 proteins (Fig 1A, S1 Table and S1 Fig). In addition, signal peptides were identified in five TvNlpC/P60 proteins (S1 Table and S1 Fig). These nine proteins are split in two distinct orthologous groups in TrichDB of four and five sequences, which we named A (NlpC_A1-4) and B (NlpC_B1-5) for simplicity, respectively (S1 Table and Fig 1). The N-termini of NlpC_A1-4 and NlpC_B1-5 are more similar to their respective orthologous group members and the level of similarity within each group also suggests that all nine TvNlpC/P60 proteins have functional signal peptides although some are not recognised as such by current bioinformatics tools (Fig 1B). In contrast all sampled animal and fungal NlpC/P60 sequences were inferred to possess a signal peptide, likely explained by the use of restricted, and taxonomically biased, reference sequences of experimentally confirmed signal peptides to train the signal peptide identifying software [30](S1 Fig.). Consistent with two orthologous groups, distinct taxonomic reports for the top 100 BlastP hits against the non-redundant protein database at the NCBI were obtained with respective members of the two groups used as query (S2 and S3 Tables), with these hits mainly derived from bacterial members of the Firmicutes and Actinobacteria. The high alien index (AI) values calculated from Blast hit lists (AI > 13, see Material and Methods, S1 Table, [31]) for all nine proteins (AI value range: 13.8-29.6) was consistent with the hypothesis that TvNlpC/P60 genes were acquired from bacteria by LGT [14,18]. A few proteins from phages infecting *Clostridium* species were also observed in the BlastP hit list (S2 Table, S3 Table). More sensitive Delta-Blast searches restricted to non-redundant proteins derived from eukaryotes led to a small cohort of hits from protists, Fungi and animals (S4 Tables). No hits were observed for the trichomonad *Tritrichomonas foetus* [32], suggesting that the LGT(s) in *T. vaginalis* were experienced by an ancestor of *T. vaginalis* following the speciation event that separated the *Tritrichomonas* and *Trichomonas* lineages [13].

**Fig 1.**
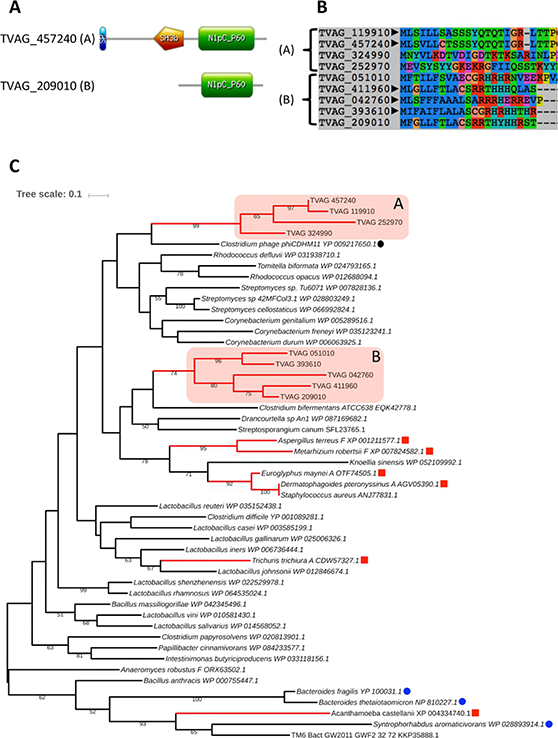
Bioinformatic analyses of the *T. vaginalis* NlpC/P60 proteins. **(A)** Structural organisation for one representative (locus tags are indicated) of each of the two clusters (A and B) of *T. vaginalis* NlpC/P60 proteins (see S1 Table and main text). Two TvNlpC/P60 proteins (cluster A), named NlpC_A1 (TVAG_119910) and NlpC_A2 (TVAG_457240, illustrated), possess a bacterial SH3 domain (orange pentagon) identified in addition to the NlpC/P60 domain (green rectangle). **(B)** N-terminus protein alignment for all nine *T. vaginalis* NlpC/P60 proteins. Locus tags are indicated and the arrowheads indicate the five entries with inferred Signal Peptide (S1 Table). Members of cluster A are more similar to each other then they are to members of cluster B and vice versa. **(C)** Phylogenetic relationship inferred by maximum likelihood from a protein alignment of 104 residues with the model LG+G4+I. Bootstrap support values above 50% are indicated and the scale bar show the inferred number of substitutions per site. The two TvNlpC/P60 proteins clusters were recovered as distinct clans (red boxes A and B). Eukaryotic sequences from non-Trichomonas species are indicated by red squares and branches. Blue and black circles are from three Gram-negative bacteria and from a phage respectively. The sequences RefSeq accession numbers are indicated. Among eukaryotes the letter “A” and “F” following the species names refers to Animal (Metazoan) and Fungi respectively.

To investigate the phylogenetic relationship between TvNlpC/P60 proteins and their homologues, all nine sequences were aligned with a selection of sequences identified from the various BlastP searches, including searches against annotated proteins derived from metagenome datasets. Metagenome-derived proteins were of particular interest as LGT in *T. vaginalis* have been characterized by a bias towards donor bacterial lineages overrepresented in mammalian gut microbiota [18]. However, none of the most similar entries to the TvNlpC/P60 proteins annotated in metagenome datasets, including from the human gut, contributed at resolving the phylogenetic position of the parasite entries (data not shown), hence these were not included in the final phylogeny. One gene derived from a ground water metagenome [33], member of the major bacterial TM6 lineage (accession KKP35888.1), was included as it clustered within a well-supported clan that included the sequence from the free-living protist *Acantamoeba castellanii*, a published candidate LGT for this species [34] (Fig 1). Additional published NlpC/P60 candidate LGTs in eukaryotes were also considered including genes from Fungi and mites [35], providing a far more appropriate phylogenetic framework to investigate the tempo and mode of LGT for all nine TvNlpC/P60 sequences compared to published analyses, which only considered one TvNlpC/P60 sequence and bacterial sequences [14,18].

A total of 50 NlpC/P60 sequences were aligned and a conservative alignment restricted to the well-conserved NlpC/P60 domain [28] was subjected to a combination of complementary phylogenetic analyses (Fig 1C and S5 Table). In contrast to a relatively recent case of LGT reported in *T. vaginalis*, which does not apparently encode any functional gene and that was possibly derived from a bacterium of the urogenital tract of the genus *Peptoniphilus* [15,36], none of the members of the vaginal microbiota included in the phylogenetic analysis (e.g. *Lactobacillus iners* and *Corynebacterium genitalium*) preferentially clustered with any of the TvNlpC/P60 sequences.

All unconstrained maximum likelihood phylogenetic analyses, including amino acid composition homogenous and composition mixture models (see Material and Methods section) recovered the members of the two TvNlpC/P60 groups in different clans (S5 Table), consistent with two distinct LGT events and with these proteins being members of different orthologous groups (S1 Table). The dissimilar indels (insert/deletion) profile among the nine aligned NlpC/P60 sequences are also consistent with two distinct groups of sequences originating from two distinct LGT events for these genes, which was followed by gene duplication events forming two distinct protein families of four (clan A) and five (clan B) TvNlpC/P60 proteins (Fig 1C and S1 Fig). However, weak bootstrap support (values <50%) for a number of stem branches in the unconstrained phylogenetic analyses did not conclusively support two independent LGT events (Fig 1C).

To further test the robustness of the distinct candidate LGT events into eukaryotes supported by all unconstrained maximum likelihood phylogenies, including the potentially two independent LGTs into *T. vaginalis*, a selection of constrained phylogenetic analyses were performed in combination with tree topology tests (S5 Table). A potential single LGT event into *T. vaginalis*, followed by gene duplications, for all nine TvNlpC/P60 genes could not be rejected with the protein single rate matrix based model (LG+G4+I) and the composition mixture model LG4X but was rejected by the more complex composition mixture model (C20 based model) (S5 Table). In contrast, all the tested hypotheses in which *T. vaginalis* might have shared the NlpC/P60 LGT event with one other eukaryotic lineage included in our analyses (different combinations of *T. vaginalis* sequences constrained to be monophyletic with a selection of the different distinct eukaryotic sequences) were rejected by all considered models, in line with the moderate to high (~70-90%) bootstrap values supporting the scattered distribution of eukaryotic sequences among bacterial homologues in the phylogeny (S5 Table). These analyses are consistent with multiple independent NlpC/P60 bacteria gene acquisition events among the sampled eukaryotes.

### The three-dimensional structure of *Trichomonas vaginalis* NlpC_A1 and NlpC_A2

To investigate the structure and function of the *T. vaginalis* NlpC proteins, we expressed and purified the proteins and undertook crystallisation experiments towards determining the structure by X-ray crystallography. Crystals were obtained for NlpC_A1 (TVAG_119910) and NlpC_A2 (TVAG_457240) using the sitting drop vapour diffusion technique. Crystals of NlpC_A1 diffracted to 1.2 Å resolution and belonged to the spacegroup P21221. Crystals of NlpC_A2 diffracted to 2.3 Å and belonged to the spacegroup P1 (S6 Table).

The structure of NlpC_A1 (PDBid 6BIM/6BIO) was determined by single isomorphous replacement using SeMet substituted protein. A single molecule is present in the asymmetric unit and all residues, including three residues that remain after cleavage of the purification tag, are visible in the electron density map. The structure of NlpC_A2 (PDBid: 6BIQ) was subsequently determined by molecular replacement using the structure of NlpC_A1 as a search model. Four copies of NlpC_A2 are present in the asymmetric unit with residues 11-275 visible in the electron density maps.

NlpC_A1 and NlpC_A2 consist of three domains (Fig 2A). The NlpC/P60 domain is located at the C-terminus of the protein and is preceded by two bacterial SH3 domains. As expected, due to the high sequence identity (90.2% identical, 27 differences, 96.4% similar), both structures are essentially identical (Fig 2B) with an RMSD on all atoms of 0.491Å (1379 equivalent atoms). For brevity, we will describe the structure for NlpC_A1 and highlight differences with NlpC_A2. The SH3b domains consist of 6 β-strands arranged in a beta barrel formation. They are joined by a shared β-strand (β7) and pack against the NlpC domain in a triangular arrangement. A short linker of approximately 10 residues joins the second SH3b domain to the NlpC domain. The NlpC/P60 domain adopts the classical papain-like fold [37] with a central β-sheet of 5 strands displaying an anti-parallel arrangement. Three helices (α2, α3, and α4) are packed against one side of the sheet and form the interface to the SH3 domains. The similarity to other NlpC proteins allows us to assign the active site residues of Cys179-His234-His246, also present in all 9 *T. vaginalis* NlpC sequences (S1 Fig). The active-site cysteine is located at the N-terminus of helix α3 while the two histidines are present in strands β15 and β16.

**Fig 2.**
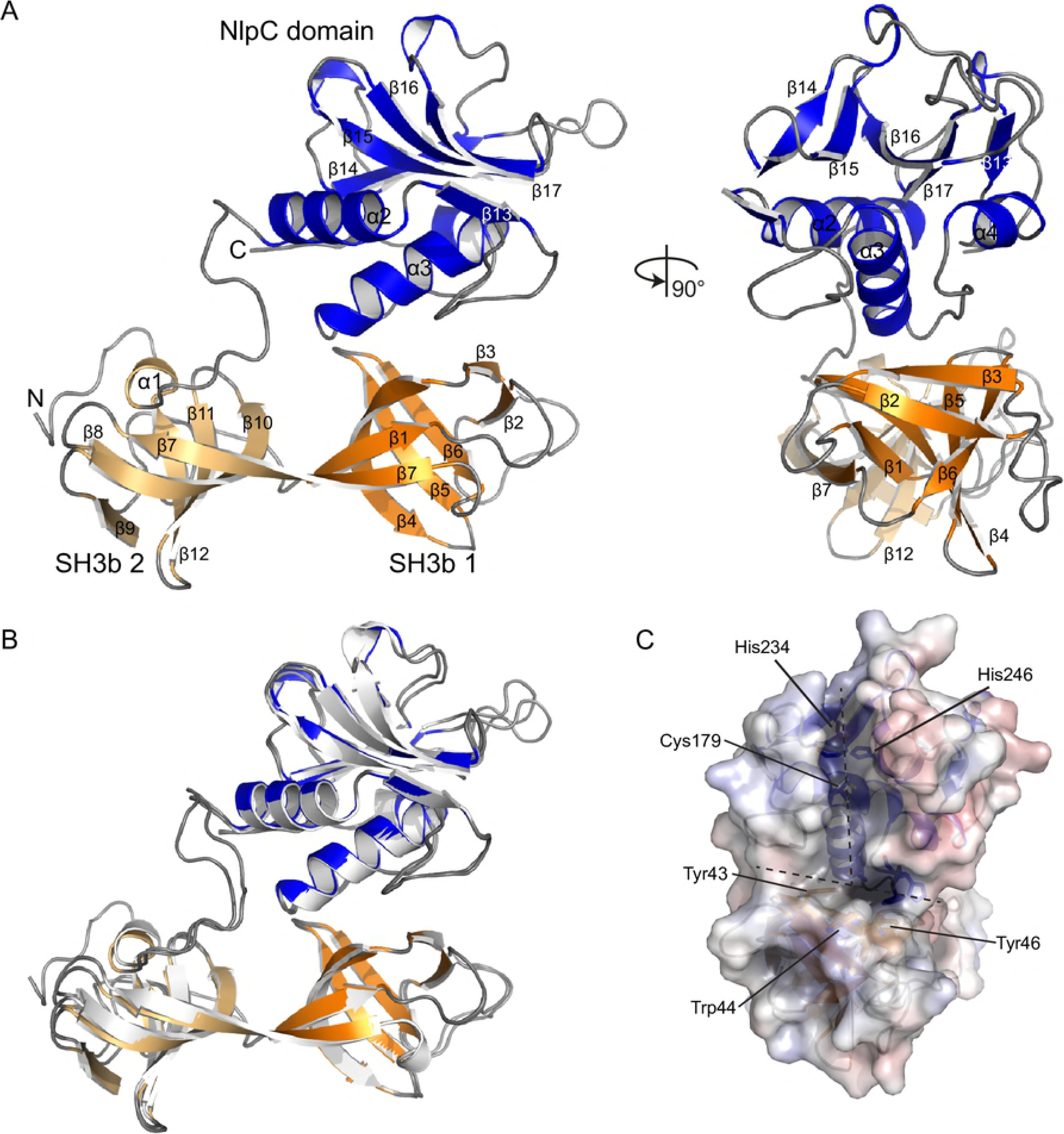
Structure of NlpC_A1 and NlpC_A2. (**A**) Orthogonal views of a cartoon representation of the structure of NlpC_A1. Secondary structure elements are labelled, N- and C- termini are indicated. The NlpC domain is colored in blue with the two SH3b domains shaded in orange. (**B**) Superposition of the NlpC_A1 and NlpC_A2 structures. The two structures superimpose with an RMSD of 0.491Å across all equivalent atoms. (**C**) Electrostatic surface representation of the NlpC_A1 active site groove including the description of the catalytic triad Cys179-His234-His246. Location of the “T” shaped groove is marked with a dashed line.

The active site residues sit in a ‘T’-shaped groove that is bounded by strands β14 and β15 in the NlpC domain, with the cross of the T formed by the strand β3 in the first SH3b domain (Fig 2C). The groove is open and lacking obvious regulatory elements that are often present in bacterial endogenous peptidases [38]. This is suggestive of both NlpC_A1 and NlpC_A2 being unregulated enzymes capable of degrading the cell-wall of bacteria.

A surface electrostatic calculation [39] demonstrates the active site groove to carry a slight positive charge with two pronounced areas of strong positive charge. The first of these is adjacent to the active site cysteine. The second is at the base of a small pocket at the interface between the NlpC domain and the first SH3b domain bounded by the residues His195, Tyr43, Trp44, Tyr46, Phe193, Gln184 (Fig 2C).

A single residue difference in NlpC_A2 is located in the vicinity of the active site where Leu169 in NlpC_A1 is Trp169 in NlpC_A2. The tryptophan residue narrows the groove around the active site cysteine in NlpC_A2. The remaining sequence differences are spread across the surface of the protein with no obvious cluster of residues that might influence function.

The presence of multiple SH3b domains, while not unusual for NlpC proteins [28], is intriguing. NlpC domains are often accompanied by accessory domains that alter substrate specificity. Previous studies of multi-domain NlpC/P60 proteins have shown that the accessory domains can exist in a flexible conformation for recognition of PG [40]. Consequently, small angle X-ray scattering (SAXS) was used to confirm the arrangement of domains within the crystal. Both NlpC_A1 and NlpC_A2 eluted as a single peak on size-exclusion chromatography and multi-angle light scattering analysis demonstrating both proteins to be monomeric in solution (S3 Fig). Comparison of the theoretical scattering from the structural models with SAXS curves is consistent with the arrangement of domains present in the crystal. This demonstrates that the NlpC domain and SH3b domains pack in a concerted arrangement in solution (S2 Fig and S7 Table).

To further investigate the role of the SH3b domains in NlpC_A1 and NlpC_A2, we undertook a series of structural similarity searches using the SSM algorithm [41]. These searches failed to identify other structures with a similar global domain arrangement. Searches using the NlpC domain only identified several other members of the NlpC superfamily that contain SH3b domains. The NlpC/P60 protein from *Bacillus cereus* (YkfC; PDBid 3H41) shares the same domain composition with two SH3b domains at the N-terminus of the protein followed by the C-terminal NlpC domain. However, alignment of the structures based upon the NlpC domains (258 atoms, RMSD 0.548 Å) reveals an alternative arrangement of the SH3b domains relative to the NlpC. Additionally, the first SH3b domain of YfkC has a large insertion consisting of three α-helices (~40 residues). This results in the active site of YkfC being more closed (typical of a recycling enzyme [42]) than those of NlpC_A1 and NlpC_A2.

### NlpC_A1 and NlpC_A2 degrade peptidoglycan

The structural comparison of NlpC_A1 and NlpC_A2 suggests that these proteins may indeed be involved in the degradation of bacterial PG and, possibly, have redundant function. To investigate this, we examined the activity of the recombinant enzymes against PG from *E. coli* as a model substrate. We used an equal mix of PG from strains MC1061 (laboratory strain, rich in tetrapeptides) and CS703-1 (carboxypeptidase mutant, rich in pentapeptides) [43,44] resulting in a substrate that contains a mixture of peptide lengths. The PG mix was incubated with NlpC_A1, NlpC_A2 or the corresponding predicted inactive versions in which the catalytic Cys residue was replaced by either Ala or Ser, followed by digestion with the muramidase cellosyl and HPLC analysis of the resulting muropeptides (Fig 3 and S3 Fig).

**Fig 3.**
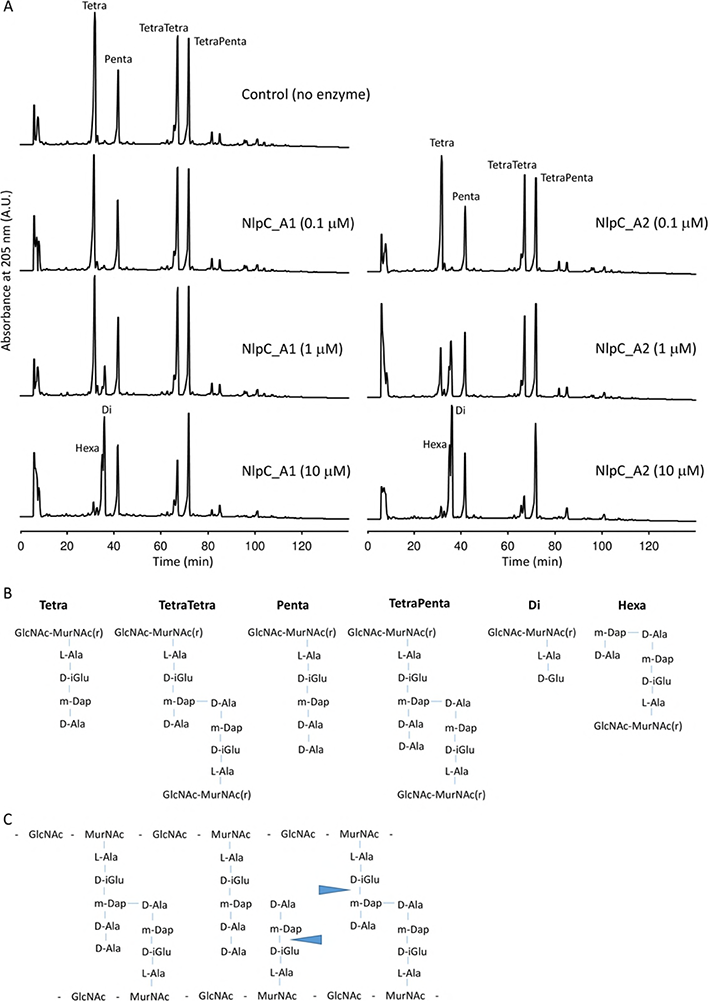
NlpC_A1 and NlpC_A2 are DL-endopeptidases. (**A**) HPLC chromatograms of *E. coli* peptidoglycan cleavage assays with control (no enzyme) and increasing concentrations of NlpC_A1 or NlpC_A2. Peaks of the major muropeptides were assigned by comparison with published literature and are labelled as Tetra, Penta, TetraTetra and TetraPenta. NlpC_A1 and NlpC_A2 produce the muropeptides Di and Hexa. (**B**) Structure of the muropeptides labelled on the chromatograms. MurNAc(r), reduced N-acetylmuramic acid; GlcNAc, N-acetylglucosamine; L-Ala, L-alanine; D-iGlu, iso-D-glutamic acid; m-Dap, meso-diaminopimelic acid; D-Ala, D-alanine. (**C**) Schematic diagram of the undigested *E. coli* peptidoglycan with the indication of the cleavage sites of NlpC_A1 and NlpC_A2 (blue arrows).

The control sample shows the retention times for the peaks corresponding to the major monomeric (Tetra and Penta) and dimeric (TetraTetra and TetraPenta) muropeptides (Figs 3A and 3B). In the presence of increasing concentration of NlpC_A1 or NlpC_A2, the Tetra peak decreased and new peaks corresponding to cleavage products appeared indicating activity against PG (Fig 3A). These new peaks resulted from the cleavage of the bond between D-isoGlu and *m-*DAP residues (Fig 3C) and classify both NlpC_A1 and NlpC_A2 as DL-endopeptidases. At the highest concentration of the enzymes the dimeric TetraTetra was digested while the muropeptides with pentapeptides (Penta and TetraPenta) were largely inert. Hence, we concluded that NlpC_A1 and NlpC_A2 are DL-endopeptidases with specificity for the tetrapeptides in peptidoglycan and with greater activity towards monomeric muropeptides. Finally, as expected, mutation of the catalytic Cys179 to either Ser or Ala in NlpC_A1 and NlpC_A2 completely abolished their activity towards PG (S3 Fig).

### The expression of endogenous NlpC_A1 and NlpC_A2 genes in *Trichomonas vaginalis*

The structure and the activity assays of NlpC proteins from *T. vaginalis* demonstrated that these LGT-acquired enzymes are capable of degrading bacterial PG. As in recent examples of bacterial-derived PG-degrading enzymes in eukaryotes [25,26], this function has been preserved in *T. vaginalis* after LGT and apparent gene duplications. To examine the potential role of NlpC enzymes in this parasite, we firstly investigated the effect of co-incubating parasite and bacteria and examined changes in the expression levels of the endogenous *nlpC* genes by RT-qPCR. Simultaneously, the mixed cultures were also plated on agar to examine the effect on bacterial cell numbers.

Bacteria and parasites were co-incubated at a 1:10 ratio in a minimal defined media at 37°C. Samples were taken at 2, 4 and 8 h time points for counting of colony forming units (cfu) when total RNA was also purified for the RT-qPCR analysis. Samples of bacteria alone (no parasite) and parasite alone (no bacteria) served as controls for cfu counts and RT-qPCR readouts, respectively. The cfu counts were used to calculate the bacterial survival as the ratio of cfu values from mixed versus bacteria-alone cultures.

We observed that *T. vaginalis* upregulates the expression of both NlpC_A1 and NlpC_A2 in the presence of bacteria. At the latest time point (8 h), transcription of both genes was upregulated by 8-9 fold (Fig 4). Interestingly, the increased expression of *NlpC* genes was accompanied by a ~20-30% reduction in the number of bacteria as compared to the control without parasites. This result shows that bacteria trigger upregulation on the expression of these *NlpC* genes in *T. vaginalis* and that their activity towards PG degradation, demonstrated by the previous *in vitro* experiments, may have an impact on bacterial survival.

**Fig 4.**
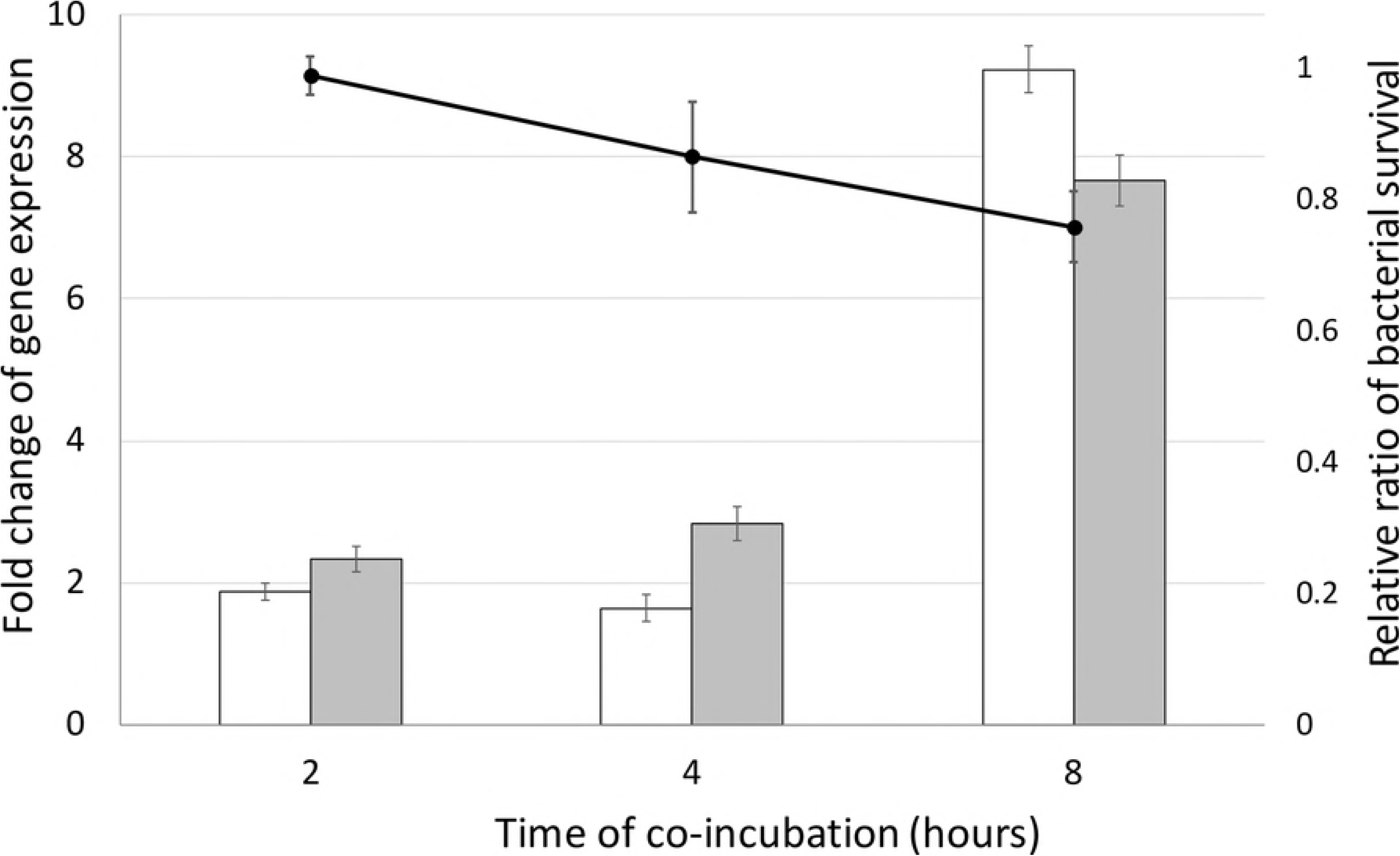
A reduction in *E. coli* numbers is accompanied by transcription upregulation of NlpC_A1 and NlpC_A2 genes in *T. vaginalis*. *E. coli* was incubated in the absence or in the presence of *T. vaginalis* at a ratio of one bacterium to ten protozoan cells and up to 8 hours. The cfu counts were used to calculate bacterial survival (line graph following the y-axis to the right). Bacterial survival was expressed as a ratio of cfu in the presence versus absence of *T. vaginalis*. Simultaneously, *T. vaginalis* in the absence of bacteria served as the base line for the RT-qPCR (bar graph following the y-axis to the left). Relative quantification of NlpC_A1 and NlpC_A2 mRNA abundance (white and grey bars, respectively) was achieved by comparing *T. vaginalis* in the presence versus absence of bacteria for each time point. The housekeeping gene HSP 70 was used as a reference and the *Ct* method was applied for relative quantification of gene expression (see Methods). The values are mean ± SD of three independent experiments.

### Subcellular localization of NlpC_A1 in *Trichomonas vaginalis*

Based on the apparent redundancy at sequence/structure (Figs 1 and 2, S1 Table), activity (Fig 3) and transcriptional regulation (Fig 4), we focused the following experiments on NlpC_A1 only. In bacteria, NlpC/P60 members often contain either a signal peptide or a transmembrane region [28]. At this stage, it was unclear if signals for subcellular compartmentalization would have been ameliorated for accurate protein targeting and sorting of the TvNlpC/P60 proteins in the parasite (S1 Table). For NlpC_A1, bioinformatics analyses identified potential signal peptides but no transmembrane regions. Hence, to ascertain the subcellular localization of NlpC_A1 in *T. vaginalis* cells, we undertook an experimental approach.

To examine this, the NlpC_A1 coding domain was expressed with a C-terminal HA-tag from a plasmid under a strong constitutive promoter. This plasmid was introduced into *T. vaginalis* by electroporation and transfected cells were drug-selected. Transfected cells expressing the HA-tagged NlpC_A1 were used in immunoassays with the anti-HA antibody. Immunofluorescence assays showed strong and consistent surface staining of transfected *T. vaginalis* cells, suggesting that HA-tagged NlpC_A1 is located on the plasma membrane (Fig 5). The immune-staining of the cell surface was patchy, reaching areas of the flagellar membrane. This was consistent with the results of a crude cell-fractionation protocol that showed the presence of the HA-tagged NlpC_A1 in the high-speed membrane pellet fraction of the Western blot (S4 Fig). Together, these findings provide experimental evidence that NlpC_A1 is likely to be located on the cell surface of *T. vaginalis*.

**Fig 5.**
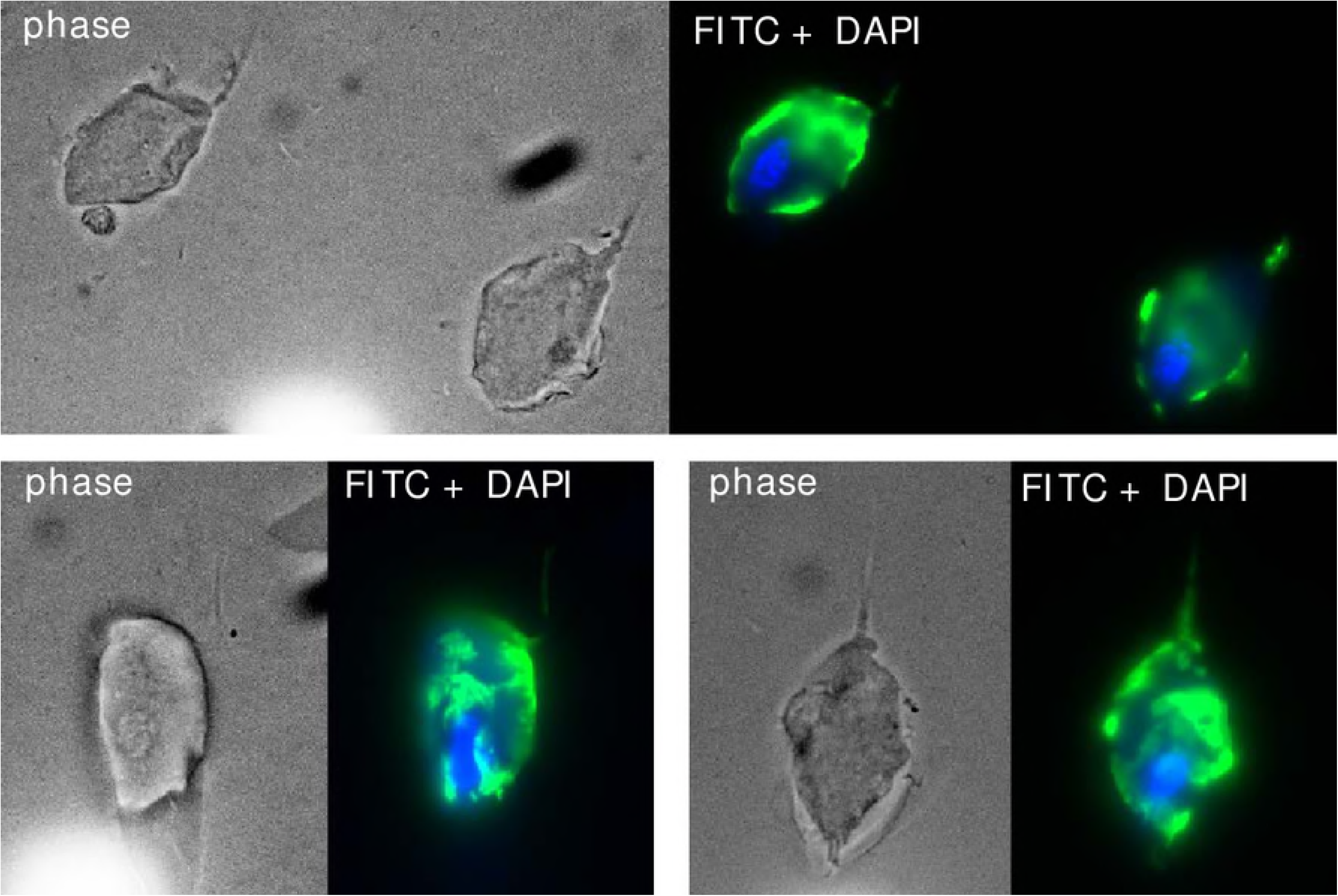
Subcellular localization of HA-tagged NlpC_A1 in *T. vaginalis*. The parasite was stably transfected with plasmids expressing HA-tagged NlpC_A1 protein. Immunofluorescence microscopy, using primary HA-specific and secondary FITC-conjugated antibodies, indicates that the NlpC_A1 is present on the membrane of the protozoan cells. The nucleus was stained with DAPI for reference. Three independent images (total magnification of 1000X) are shown.

### Profound phenotypic changes upon overexpression of exogenous NlpC_A1 gene

With the observation that *T. vaginalis* were able to reduce the population of bacteria in mixed cultures (Fig 4), we tested if overexpressing NlpC_A1 would enhance this phenotype. With the initially used cell ratio of 1:10 (bacteria:protozoan), NlpC_A1 *T. vaginalis* transfectant virtually eliminated bacteria in the co-cultures after just 1 h of co-incubation (S5 Fig). Therefore, we inverted the cell ratio to 10:1 (bacteria:protozoan) and included an empty-plasmid transfectant (expressing only the neomycin selection marker) as a negative control. In addition, we also tested if bacterial clearance depends on the specific activity of NlpC_A1 with the expectation that mutation of a critical catalytic residue (C179S) should diminish this phenotype. For each time point, bacterial cfu numbers were determined by spotting and spreading co-cultures on agar plates. The obtained data were reported as the relative cfu obtained from mixed cultures containing *T. vaginalis* transfected with either WT NlpC_A1 or the mutant NlpC_A1 versus the empty-plasmid *T. vaginalis* transfectant (Fig 6).

**Fig 6.**
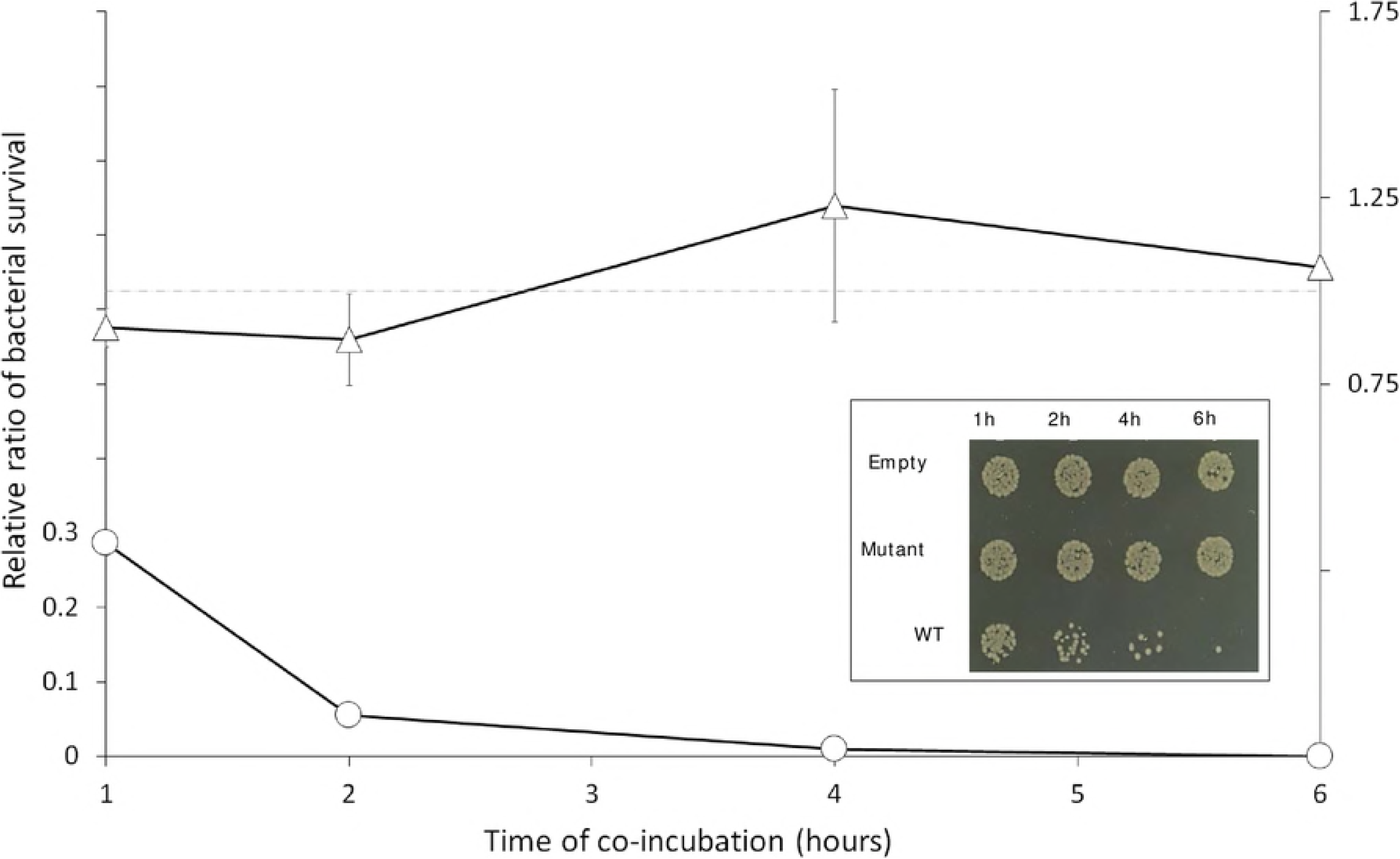
*T. vaginalis* promotes clearance of *E. coli* DH5α from mixed cultures when over-expressing NlpC_A1 constitutively. *T. vaginalis* was stably transfected with plasmids expressing no NlpC_A1 protein (Empty), the HA-tagged NlpC_A1 mutated at the catalytic residue C179S (Mutant) or the HA-tagged NlpC_A1 wild-type (WT). *E. coli* was incubated in the presence of each of the transfected *T. vaginalis* at a ratio of ten bacteria to one protozoan cell and up to 6 hours. *E. coli* cfu counts were used to determine bacterial survival in the presence of *T. vaginalis* expressing mutant (triangles) or WT (circles) NlpC_A1. The inset figure illustrates the growth of *E. coli* when co-incubated with each of the *T. vaginalis* transfectants and at each time point, as indicated. Undiluted mixed cultures were individually spotted on a LB-agar plate. The values are mean ± SD of three independent experiments.

We observed a significant drop in bacterial cfu counts when *T. vaginalis* overexpresses the wild-type NlpC_A1 exogenously (Fig 6). This effect could be readily observed on the spot plates during the course of incubation (Fig 6, inset) where a dramatic and time-dependent reduction on bacterial numbers was noticeable on the spots coming from mixed cultures containing *T. vaginalis* transfected with the WT NlpC_A1. By contrast, bacteria maintained similar viability over the course of the experiment in mixed cultures containing *T. vaginalis* transfected with either the inactive C179S mutant NlpC_A1 or the empty plasmid (Fig 6, inset). Quantification of cfu confirmed the drastic loss in bacterial viability when *T. vaginalis* overexpressed the wild-type, active form of NlpC_A1 (Fig 6, line graph). After one hour of co-incubation, the bacterial population was reduced by 70% and bacteria were virtually eliminated after 4-6 hours. *T. vaginalis* transfected with the catalytic-inactive NlpC_A1 (C179S mutant) or empty plasmid behaved similarly, resulting in a constant bacterial cfu ratio of ~1 (dotted line) throughout the course of the experiment.

To further support the observation that the activity of NlpC_A1 is responsible for this phenotype, *T. vaginalis* transfectants were incubated with the *E. coli* strain CS703-1 which contains a PG highly enriched for pentapeptides [44]. We observed previously that the recombinant NlpC_A1 was incapable of digesting pentapeptides in PG (Fig 3). Therefore, the ability of bacterial clearance by *T. vaginalis* overexpressing the wild-type NlpC_A1 might be compromised when co-incubated with bacteria carrying a pentapeptide-rich PG (CS703-1) as compared to bacteria carrying a tetrapeptide-rich PG (DH5α) (Fig 7). Visual inspection of the undiluted mixed cultures on spread plates after 1 h of co-incubation showed a significant reduction on the numbers of bacterial colonies regardless of the strain, CS703-1 or DH5α, when *T. vaginalis* overexpresses the wild-type form of the NlpC_A1 (Fig 7, left). However, quantification of cfu indicated that *T. vaginalis* overexpressing the wild-type NlpC_A1 was partially impaired from clearing CS703-1 as compared to DH5α by a factor of ~10 (Fig 7, right). This result is consistent with the biochemical evidence that this enzyme is unable to digest pentapeptides in PG.

**Fig 7.**
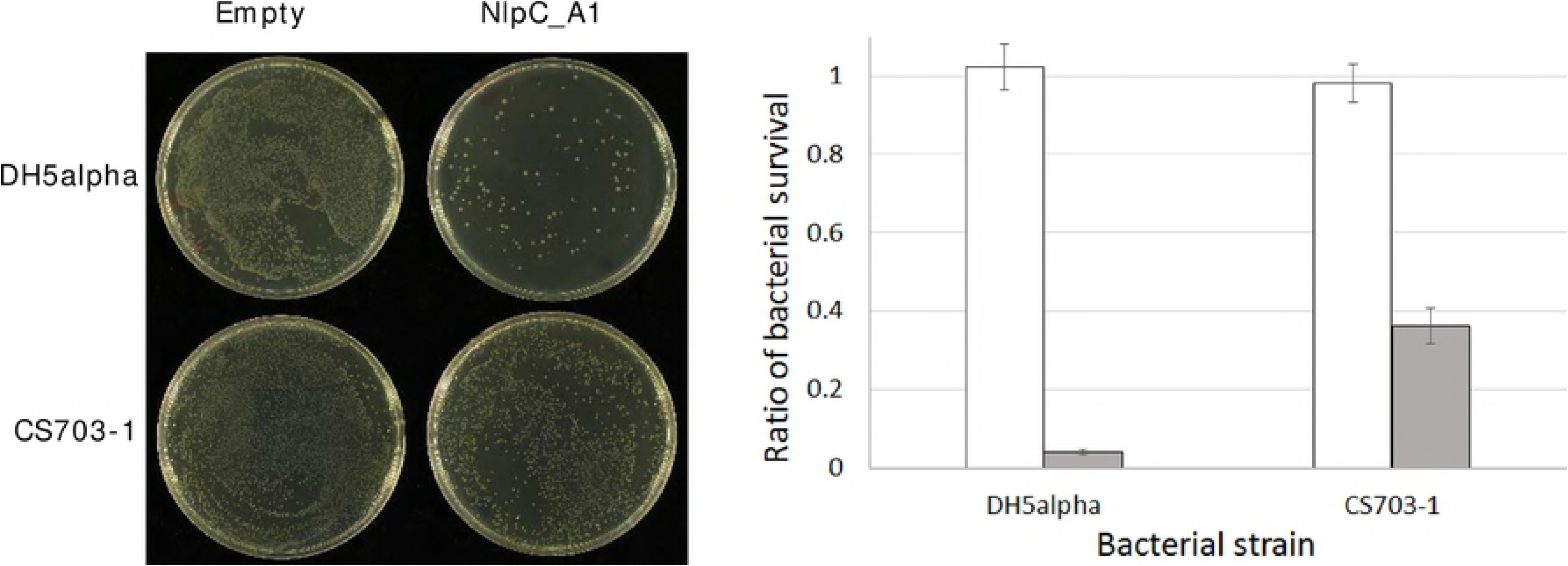
*T. vaginalis* over-expressing NlpC_A1 is partially impaired on clearing up a pentapeptide-rich peptidoglycan bacteria from mixed cultures. *T. vaginalis* was stably transfected with plasmids expressing no NlpC_A1 protein (Empty) and HA-tagged NlpC_A1 wild-type (NlpC_A1). *E. coli*, strains DH5α and CS703-1, were incubated alone or in the presence of each of the transfected *T. vaginalis* at a ratio of four bacteria to one protozoan cell for one hour. **Left**. To illustrate the bacterial growth inhibitory effect of *T. vaginalis* Empty versus WT against these strains of *E. coli*, undiluted mixed cultures were plated on LB-agar. **Right**. To measure the levels of bacterial survival, dilutions of the mixed cultures were plated on LB-agar and cfu counts were obtained. For each *E. coli* strain (DH5α and CS703-1), cfu counts were used to calculate bacterial survival in the presence of the Empty *T. vaginalis* (white bar) or *T. vaginalis* expressing NlpC_A1 wild-type (grey bar).

## Discussion

To our knowledge, this is the first study to reveal that a laterally-acquired group of genes enables a eukaryotic mucosal pathogen to control bacterial population. Our combined unconstrained and constrained phylogenetic analyses support the hypothesis that these genes were acquired from different bacteria, or possibly one bacteria and a phage, and expanded through subsequent gene duplications during evolution of the *Trichomonas* lineage. However, we could not establish with precision the likely bacterial donor lineage from our phylogenetic analyses, either because genomes of bacteria closely related to the donor(s) have not yet been sequenced and/or because the gene transfers took place in the distant past making it more difficult to infer evolutionary relationships due to important sequence divergence eroding phylogenetic signal. While some phages encode candidate or functional NlpC/P60 cell wall peptidases [28], a possible phage origin for TvNlpC/P60 members of clan A was only weakly supported in all of our unconstrained phylogenetic analyses (Fig. 1C), The combination of BlastP searches, unconstrained and constrained phylogenetic analyses indicates that the two TvNlpC/P60 distinct gene families were likely acquired by two independent LGTs and further supports several independent LGT events in other eukaryotic lineages, as recently reviewed by Husnik and McCutcheon [22].

Consistent with relatively ancient LGT event(s) into the *Trichomonas* lineage, analyses of GC% profile variation of the nine scaffolds encoding the nine TvNlpC/P60 genes, we observed that seven genes were located in segments with average GC% composition characteristic of the *T. vaginalis* genome (~31-32 GC%) and were not located close to segmentation points (S1 Table and S6 Fig). This is in contrast to the relatively recent candidate LGT likely derived from a bacterial donor from the urogenital tract identified on a ~53 kbp *T. vaginalis* genome sequence scaffold [35]. A segment of ~34 kbp encodes 27 proteins highly similar (range: 79–98% identity) and with long stretches of gene synteny with the bacteria *Peptoniphilus harei*. This case is characterised by a strong GC% segmentation point between the ~34 kbp segment encoding the 27 bacterial-like genes and the rest of the scaffold [15,36]. One TvNlpC/P60 gene (TVAG_393610, DS113268, S1 Table) was located beside a segmentation point on a region with >30 GC% content which was correlated with the presence of repetitive elements, well known to litter the highly repetitive genome of *T. vaginalis* [14] rather than a Bacterial-like segment of DNA. By investigating the GC% profile of the open reading frames (ORFs) from all nine TvNlpC/P60 genes, a correlation between codon GC% profile and clan membership was observed. The ORFs from the respective members of clan A were more similar to each other (GC% from 38.5 to 40.6) than to the ORFs from the other clan B (GC% from 44.1 to 50.0) (S1 Table). This is also consistent with two genes families derived from distinct LGT events and followed by gene duplications.

Transcriptional upregulation of NlpC_A1 and NlpC_A2, as induced by glucose starvation [16], and by bacteria (as shown here), supports the functional integration of these genes of bacterial origin in *T. vaginalis*. These independent observations indicate that these genes have acquired optimal codons and promoters that confer proper gene expression and regulation upon specific environmental conditions. In other words, these genes have been fully integrated into the biology of the parasite through amelioration [45]. As anticipated for NlpC/P60 enzymes, the tri-dimensional structure and catalytic function have been preserved in the two members of this group of enzymes (NplC_A1 and NlpC_A2) that we have characterized here. These LGT-derived genes in *T. vaginalis* encode functional DL-endopeptidases that have specific activity towards PG, a unique polymer that constitutes an essential component of the bacterial cell envelope.

NlpC_A1 and NlpC_A2 are apparently redundant in structure and function. Their high-resolution structures revealed a classical papain-like NlpC/P60 domain in addition to two bacterial SH3 domains under a specific arrangement. The open ‘T’-shaped groove includes the catalytic triad Cys-His-His which was shown here, as predicted, to be necessary for its activity. Both enzymes are active against PG cleaving the chemical bond between D-isoGlu and *m*-DAP residues on tetrapeptides in monomeric and dimeric peptidoglycan subunits. However, curiously, neither of them can cleave the same bond if the PG substrate is a pentapeptide or tetrapentapeptide. During bacterial cell wall synthesis, the fifth (terminal) D-alanine residue of pentapeptide donor peptides is removed during transpeptidation reactions, but newly made (nascent) peptidoglycan still contains a high percentage of pentapeptides in monomers and dimers [46]. In many species, for example *E. coli*, the majority of these pentapeptides are trimmed to tetrapeptides by DD-carboxypeptidases during peptidoglycan maturation, resulting in a mature, tetrapeptide-rich peptidoglycan [47]. Therefore, *T. vaginalis* NlpC_A1 and NlpC_A2 are apparently optimized to digest mature bacterial cell walls consistent with functional enzymes targeting mature bacterial substrates in the native habitat of the parasite.

We found that, when bacteria and parasites were co-incubated in a minimal media, the expression of NlpC_A1 and NlpC_A2 was upregulated and this was accompanied by a reduction on the numbers of viable bacteria. Interestingly, glucose restriction also leads to transcriptional upregulation of these two *T. vaginalis* NlpC/P60 genes [16]. In the vaginal microenvironment, where glucose can be restricted and bacteria are abundant, it is possible that *T. vaginalis* overexpresses these enzymes to target bacteria as a source of nutrition. PG turnover during bacteria replication or bacterial lysis will release PG fragments among other metabolites which *T. vaginalis* could scavenge for, by using its own repertoire of NlpC/P60 and perhaps additional enzymes.

In bacteria, NlpC/P60 enzymes are often secreted into the periplasm or growth medium. To date, *T. vaginalis* NlpC/P60 enzymes have not been detected in proteomics surveys of surface or secreted proteins [48–50], perhaps because their expression depends substantially on environmental factors such as glucose restriction [16] and presence of bacteria (as found in this study). In *T. vaginalis*, we have shown that NlpC_A1 localizes on the surface of the parasite. However, the process by which these enzymes reach the peptidoglycan of the target bacteria remains to be determined. In the case of Dae2 (a NlpC/60 protein), which was laterally inherited by the tick *Ixodes scapularis* from bacteria, bacteriolytic activity is only detected if this enzyme is delivered to the periplasm of *E. coli* or if the outer membrane is permeabilized [26].

Although the overexpression of NlpC_A1 in *T. vaginalis* significantly enhances the ability of the pathogen to control bacterial population in co-cultures, the purified recombinant enzyme is not itself bacteriolytic. This was demonstrated by incubating *E. coli* with the purified enzyme and comparing to the well-known bacteriolytic activity of lysozyme (S7 Fig). Therefore, other factors or additional enzymes such as cationic antimicrobial peptides and lysozyme [51] (of either microbial or human origins) might possibly enable *T. vaginalis* NlpCs to reach the peptidoglycan of vaginal bacteria. Vaginal secretions are known to have the highest level of lysozyme among other human mucosa-derived fluids [52]. Finally, direct contact between parasite and bacteria and parasite-mediated phagocytosis could possibly facilitate the action of these enzymes.

The profound reduction in viable bacteria, when a functional exogenous copy of NlpC_A1 gene was constitutively expressed under a strong promoter in *T. vaginalis*, provided *T. vaginalis* a striking ability of controlling bacterial population in co-cultures even in the presence of a 10-fold excess of bacteria. This phenotype was dependent on the activity of this enzyme since *T. vaginalis* transfected with the catalytic-inactive NlpC_A1 (C179S) did not display such activity beyond the background. Additionally, this phenotype replicates the inability of digesting the pentapetide-rich PG as observed with the recombinant enzyme. We have not characterized all members of this family in *T. vaginalis* neither examined their broad bacterial activity yet. However, our study suggests that *T. vaginalis* NlpC enzymes may act as cell-wall degrading toxins. This is supported by their structural data, cytolocalization and the impact on bacterial survival from mixed cultures when the parasite overexpresses NlpC_A1. In contrast to the housekeeping NlpC/P60 PG hydrolases of bacteria, the structures of *T. vaginalis* NlpC_A1 and NlpC_A2 revealed an open and highly accessible active site which is characteristic of a cell-wall degrading toxin [53].

The functional redundancy of *T. vaginalis* NlpC_A1 and NlpC_A2 may be complemented by the other NlpC/P60 members in the parasite. Such apparent redundancy in peptidoglycan enzymes is well known in the PG biogenesis of bacteria and mostly evident in PG hydrolysis. *E. coli*, for instance, has 6 PG endopeptidases, 4 amidases and 8 lytic transglycosylases with distinct biochemical properties [54]. Together, the combined activities in each set of seemingly redundant enzymes allow robust growth even at different conditions. This is important because the properties of the periplasm (e.g. osmolarity and pH), where these enzymes function, depends on the environmental conditions [54]. Similarly, *T. vaginalis* could maintain a seemingly redundant set of NlpC/P60 enzymes to target PG of different vaginal bacteria (e.g. Gram-negative and/or –positive), at different conditions and locations. Indeed, these different enzymes may act inside phagosomes or extracellularly including at the cell surface or in secretions. Consistent with these possibilities, one gene from clan B (TVAG_209010) was characterised by a distinct higher level of transcription in two RNASeq surveys compared to all other TvNlpC/P60 genes [16,17] with in particular a dramatic upregulation across 36 hours of incubation in the absence of glucose [16] (S1 Table).

At this stage, the contribution of these enzymes to pathogenesis is still unknown. However, several possibilities can be envisaged. Firstly, *T. vaginalis* might benefit from the released PG fragments and other bacteria derived metabolites by using them as nutrients. Secondly, controlling bacterial populations will not only reduce microbial competition but also prevent the protective function of the host microbiota favouring *T. vaginalis* propagation. Thirdly, molecules from PG degradation are known ligands of pattern recognition receptors in the host [55,56]. Therefore, *T. vaginalis* NlpC activity may have a role on modulating immune responses altering the outcomes of the infection. Lastly and more speculative, these enzymes may explain the association of *T. vaginalis* with a particular community of vaginal bacteria [12] and/or with mycoplasmas [57]. Mycoplasmas are very frequently found as intracellular symbionts of *T. vaginalis*. We speculate that the lack of PG and a rigid cell wall may have allowed them to become successful symbionts of this parasite.

The interaction of *T. vaginalis* with the microbiota has historically escaped much consideration from parasitologists, microbiologists and clinicians alike. Our study reports a novel aspect on the biology of *T. vaginalis*. It indicates that this parasite utilizes hydrolases active against peptidoglycan, a major and very specific component of bacterial cell walls. Interestingly, our combined data support a model in which these PG DL-endopeptidase genes from *T. vaginalis* (i) were acquired by LGT from bacteria, (ii) were expanded and ameliorated through evolution and (iii) are being used by the parasite to control bacterial populations. In future, it will be important to understand the specificity, redundancy and/or complementarity of this expanded family of LGT-acquired genes with regards to their potential impact on the composition of the human vaginal microbiota and how these might contribute to pathogenesis.

## Materials and Methods

### General sequence analyses and phylogeny

The structural organization of the *T. vaginalis* NlpC/P60 proteins was investigated with InterProScan [58]. The potential presence of N-terminal signal peptides for RER targeting was inferred with Phobius [59], SignalPv4.1 [60] and SPOCTOUS [61]. In order to compile an alignment of homologues, BlastP searches at the NCBI were performed against nr, RefSeq or Env_nr (Metagenomic proteins) and with one protein member of each of the two *T. vaginalis* NlpC/P60 clusters identified at EupathDB/TrichDB [29] (S1 Table, illustrated in Panel A). In addition, a more sensitive DELTA-Blast search against RefSeq was performed to identify potential homologues in eukaryotes other than *T. vaginalis*.

To investigate the phylogenetic position of the nine *T. vaginalis* NlpC/P60 proteins, selections of top hits of the different BlastP searches were combined to generate an alignment maximising taxonomic representation. The alignment was trimmed with trimAl (option: gappyout) [62] to 104 residues to ensure that a conservative selection of well-aligned residues was used for the phylogenetic inference. The alignment was analysed with SMS [63] in order to establish the best fitting model based on single amino acid replacement matrices available within PhyML for the NlpC/P60 protein alignment, which was LG +G4 +I for both the Akaike information criterion (AIC) and Bayesian information criterion (BIC). Additional protein mixture evolutionary models were also considered (LG4X [64] and C20 model [65]) as they were shown to be more reliable in the context of alignments were sequences evolve at different rate and some aligned sites are saturated in relation to phylogenetic signal/noise. Unconstrained phylogenetic analyses were performed with either PhyML [66] within SEAVIEW v4.5.3 [67] or IQ-TREE [68,69] with either unique exchange rate matrix based models (LG+G4+I or LG+G4+I+F) or empirical protein mixture models (LG4X+R+F or C20+R). Constrained analyses forcing specific relationships including (i) all nine TvNlpC/P60 entries being monophyletic or TvNlpC/P60 being monophyletic with either (ii) two fungal (*Aspergillus* and *Metarhizium*), (iii) the amoeba (*Acanthamoeba*) or (iv) the parasitic worm (*Trichuris trichura*) sequences were performed with IQ-TREE, as were Approximate Unbiased (AU) tree topology tests that is currently the most appropriate test for comparing multiple trees [70]. The maximum likelihood tree (model LG+G4+I) was edited using iTOL [71] to generate Figure 1B.

The alien index (AI) for the TvNlpC/P60 genes were calculated as described by Rancurel et al. [31] using BlastP outputs from the NCBI Blast server (BLAST+ 2.8.0-alpha released at: https://blast.ncbi.nlm.nih.gov/Blast.cgi) and recording the E-values of the top Prokaryotic hit (in all cases a Bacterium) and the top non-self eukaryotic hit (all values are listed in S1 Table).

The GC% variation profiles of the nine scaffolds that encode the TvNlpC/P60 genes were analysed with GC-profile server [72]. Details of the analyses for each scaffold are provided in the S6 Fig with the GC% variation profiles. The GC% of the TvNlpC/P60 ORFs were analysed with the CAIcal server [73]. All values are listed for each gene/scaffolds in S1 Table.

### Microbial cultures and co-incubation assay

*Escherichia coli* strains BL21(DE3) and DH5α (Invitrogen), MC1061 and CS703-1 [44] were grown in Luria-Bertani (LB) media with agitation at 37 °C. *Trichomonas vaginalis* reference strain G3 was cultured in TYM medium [74] supplemented with 10% horse serum, 10 U/mL penicillin and 10 ug/mL streptomycin (Invitrogen) with no agitation at 37 °C.

To measure the effects of *T. vaginalis* on the survival of *E. coli*, the following co-incubation assay protocol was developed. Prior to the assays, the number and viability of *E. coli* and *T. vaginalis* were assessed and only cultures with at least 95% viability were used. For *E. coli*, these were assessed by flow cytometry using BD^TM^ cell viability kit and the AccuriTM C6 flow cytometer (BD Biosciences) as previously described [75]. *T. vaginalis* was grown in the absence of antibiotics to a maximum concentration of ~1 X 10^6^ cells/mL, as counted by a haemocytometer, when virtually all cells were alive and motile. Microbial cultures were spun down, washed and resuspended in antibiotic- and serum-free Keratinocyte-SFM media (Gibco). *T. vaginalis* (5 × 10^5^ cells/mL) was mixed with bacteria at specified cell ratios in a 12- or 24-well tissue culture plate and in a volume of 0.5-1.0 mL respectively and at 37 °C. Cell ratio and time of incubation were indicated for each experiment. As controls, *T. vaginalis* and bacteria were incubated alone in parallel and under the same conditions. When countable numbers of bacterial colony forming units (cfu) were needed, serial dilutions of the co-cultures were done in sterile water. Undiluted or diluted co-cultures were spotted or spread-plated on LB-agar plates and incubated overnight at 37 °C. This procedure is selective for *E. coli*, preventing growth of *T. vaginalis*. Three independent experiments were carried out. Plates were photographed using a Canon 40D camera with cannon 50 mm macro f2.5 or 100 mm macro f2.8 lenses and analysed.

### Reverse transcription and quantitative PCR (RT-qPCR)

To compare the changes on expression of *T. vaginalis* NlpC_A1 and NlpC_A2 genes upon exposure to bacteria, total RNA was obtained with TRIzol (Invitrogen) from the co-incubation assay cultures containing either *T. vaginalis* with *E. coli* or *T. vaginalis* alone. This RNA was treated with DNase I (Ambion), ethanol precipitated and cleaned further with the mini RNeasy MinElute clean-up kit (Qiagen). Reverse transcription was achieved from 5 μg of this purified total RNA using Superscript RT III supermix and oligo dT primers, as recommended (Invitrogen). To ensure that RNA samples were DNA-free, reactions omitting the reverse transcriptase (-RT) were done in parallel.

Primers targeting *T. vaginalis* NlpC_A1 or TVAG_119910 (forward-TCACAATTCCAACCCAATCTG; reverse-CTCCGTCATTTGCACCATCT) and NlpC_A2 or TVAG_457240 (forward-TAAGACCAAGCTTGGCTGC; reverse-TTCCGACATACATTCCGAC) were used for quantitative real-time PCR (qPCR). Primers targeting *T. vaginalis* HSP70 TVAG_237140 (forward-ACACAGGCGAGAGACTCGTT; reverse-TCTTTGACCCAAGCATCTCC) were used for normalization. qPCR assays were carried out to validate specificity and efficiency of each pair of primers and, as recommended, the threshold cycle (Ct) and the base line were set up (7900 HT-Realtime instructions, Applied Biosystems). With this validation, the Ct method was then applied for the relative quantification of NlpC gene expression using the Ct method (equation 2^-∆∆CT^). qPCR reactions were carried out using 10 ng of cDNA, 200 nM of each primer and PowerUP SYBR Green Master Mix (Thermofisher) following recommendations (Applied Biosystems). For each one of the three independent co-incubation assays, PCR reactions were carried out in triplicates. In addition to the -RT samples, parallel reactions without any template (water instead) were added as negative controls. qPCR data for NlpC expression were normalised against HSP70 and analysed using the SDS 2.3 and RQ Manager 1.2 applications (Applied Biosystems). Subsequent statistical analyses were then carried out using the Relative Expression Software Tool (REST©) [76].

### Plasmids, PCR and DNA cloning

The MasterNeo plasmid was used for cloning and expression of NlpC_A1 in *T. vaginalis* [77]. The insertion of a coding sequence into this plasmid, using *Nde*I and *Asp*718 restriction sites, provides a strong constitutive promoter for transcription and a double-hemagglutinin (HA) tag on the C-terminus of the protein. In addition, MasterNeo can replicate in *E. coli* and *T. vaginalis* allowing selection of transformants by ampicillin and G418 respectively [77]. The full coding sequence of NlpC_A1 was PCR-amplified from *T. vaginalis* genomic DNA with specific primers carrying those restriction sites and cloned into MasterNeo. A MasterNeo plasmid containing no exogenous gene (empty-plasmid), except for the neomycin selectable marker, was kindly donated by Patricia Johnson (UCLA). Transfection of *T. vaginalis* was achieved using the GenePulser Xcell electroporator (Bio-Rad), as previously described [77].

The pET47b plasmid (Novagen) was used for cloning and expression of NlpC_A1 and NlpC_A2 in *E. coli*. Coding sequences were PCR-amplified from *T. vaginalis* genomic DNA with specific primers aiming a ligation-independent cloning strategy [78]. Transformation of *E. coli* was achieved by standard heat-shock or electroporation protocols. Site-direct mutagenesis of NlpC_A1 and NlpC_A2, a single substitution of cysteine-179 to either serine or alanine, was achieved by inverse PCR using either the MasterNeo or pET47b plasmids as DNA template. All PCR reactions were performed with high-fidelity Phusion polymerase (ThermoFisher Scientific) or KOD polymerase (Novagen) using the Mastercycler pro thermocycler (Eppendorf). All PCR-derived inserts were fully sequenced from their individual plasmids (http://www.sbs.auckland.ac.nz/en/for/researchers-2/cpgm.html).

### Expression and purification of recombinant NlpC_A1 and NlpC_A2

NlpC_A1 and NlpC_A2 were expressed in *E. coli* BL21(DE3), a strain optimized for protein expression (Invitrogen). Cultures containing 25 μg/mL kanamycin were prepared at 37 °C prior to induction of protein expression. A single colony from a fresh transformation was seeded into a 5 mL culture. Following incubation overnight, this culture was scaled up to 750 mL. Once culture reached an OD (600 nm) of 0.4-0.6, they were transferred to 18 °C for 30 min. Protein expression was then induced by the addition of 0.5 mM IPTG and further incubation at 18 °C overnight. Cells were harvested by centrifugation, resuspended in 20 mM Tris/HCl pH 7.8 and frozen until required.

To purify the proteins, cell pellets were thawed and resuspended in lysis buffer (20 mM Tris/HCl pH 7.8, 300 mM NaCl, 0.5 mM TCEP, 20 mM imidazole, 10% glycerol). To preserve the activity of the proteins interest (which are candidate /peptidases), protease inhibitors were omitted from the purification procedure. Cells were lysed using a constant systems cell disruptor and the resulting lysate cleared by centrifugation (14000 *xg*, 30 min, 4 °C). NlpC proteins were then purified by immobilised metal affinity chromatography and eluted using a step gradient in 20 mM Tris/HCl pH 7.8, 300 mM NaCl, 0.5 mM TCEP, 300mM imidazole. Fractions containing NlpC proteins were pooled and dialysed overnight against 10 mM Tris/HCl, 25 mM NaCl, 0.1 mM TCEP with the addition of 3C-protease to remove the N-terminal His-tag.

NlpC proteins were further purified by anion exchange chromatography using a 6 mL Resource Q column and eluted on a gradient of 0-0.5 M NaCl. This was followed by a final size-exclusion chromatography step using a Superdex 75 16/60 column equilibrated in 10 mM Tris/HCl pH 7.8, 150 mM NaCl, 0.1 mM TCEP and eluted isocratically. Proteins were concentrated using a 10KDa cut-off Vivaspin centrifugal concentrators. Protein concentration was determined by UV/Vis-spectroscopy using the following theoretical masses and extinction coefficients: NlpC_A1 31415 Da, ε _280nm_ 63510 M^-1^cm^-1^; NlpC_A2, 31394 Da, ε _280nm_ 66155 M^-1^cm^-1^. For the Seleno-Methionine (SeMet) substituted NlpC_A1, expression in BL21(DE3) cells was achieved in PASM-505 media using the inhibition method [79] and the recombinant protein was then purified as described above.

### Protein structure determination of NlpC_A1 and NlpC_A2

Crystallisation of NlpC_A1 (16 mg/mL) and NlpC_A2 (14.5 mg/mL) were undertaken by sitting drop vapour diffusion using 96-well intelli-plate crystallisation trays using the morpheus screen [80] and our in-house robot screens [81]. Drops consisted of 0.5 μL of protein mixed with 0.5 μL of reservoir solution. Crystals of NlpC_A1 grew in condition A1 of the Morpheus screen (10% PEG20K, 20% PEG550mme, 0.06 M Divalents, 0.1 M imidazole-MES pH 6.5). Crystals of NlpC_A2 grew in 0.2 M Ammonium Fluoride, 20% PEG3350.

For data collection, individual crystals of NlpC_A1 were harvested and directly frozen by plunging into liquid nitrogen. Crystals of NlpC_A2 were harvested and briefly transferred into a cryoprotectant solution consisting of reservoir solution supplemented with 20% glycerol before freezing in liquid nitrogen. X-ray diffraction data were collected using our in-house X-ray generator consisting of a Rigaku Micromax-007HF rotating anode equipped with a copper anode, osmic focussing optics and MAR345 detector. Crystals of NlpC_A1 were also stored and sent to the Australian Synchrotron for high-resolution data collection.

The structure of NlpC_A1 was determined by single isomorphous replacement using Native and SeMet data collected from our in-house X-ray suite. Selenium sites were identified using the SHELXC/D/E pipeline [82] within the CCP4 program suite [83]. Phases were then used to build an initial model in PHENIX [84] followed by iterative rounds of model building and refinement in Coot [85] and PHENIX [84]. The structure of NlpC_A2 was determined by molecular replacement using PHASER [86] with NlpC_A1 as a search model. The model was built and refined in Coot and PHENIX.

### Enzymatic activity of recombinant NlpC_A1 and NlpC_A2

Equal quantities of purified PG (0.5 mg/mL) from *E. coli* strains MC1061 [43] and CS703-1 [44] were mixed and used for the activity assays. NlpC_A1 or NlpC_A2 were incubated at varied concentrations (0.1, 1.0 and 10 μM) with the *E. coli* PG mixture in 20 mM Tris/HCl pH 7.5, 150 mM NaCl for 4 h at 37 °C on a Thermomixer at 750 rpm. A control sample received no enzyme. The inactive mutants NlpC_A1(C179A), NlpC_A1(C179S), NlpC_A2(C179A) and NlpC_A2(C179S) were used at the highest concentration (10 μM). The reaction was stopped by the addition of 1/4 volume of 80 mM sodium phosphate pH 4.8 and incubation at 100 °C for 5 min. The samples were incubated overnight with 10 μg of cellosyl (Hoechst, Frankfurt am Main, Germany) at 37 °C on a Thermomixer at 750 rpm. Following the second incubation, the samples were incubated at 100 °C for 10 min and centrifuged at room temperature for 15 min at 16,000×*g*. The muropeptides present in the supernatant were reduced with sodium borohydride and separated by HPLC as described [87,88].

### Subcellular localization of NlpC_A1

Prediction of subcellular localization of NlpC_A1 was attempted using SignalP4.1 Server (http://www.cbs.dtu.dk/services/SignalP/), PHOBIUS (http://phobius.sbc.su.se/), TargetP (http://www.cbs.dtu.dk/services/TargetP/) and TMHMM (http://www.cbs.dtu.dk/services/TMHMM/). Nevertheless, taking advantage of the C-terminally HA-tagged NlpC_A1 in the transfected *T. vaginalis*, subcellular localization was verified experimentally by immunofluorescence microscopy, cell fractionation and Western blot.

A standard immunofluorescence protocol was used for localization of the HA-tagged NlpC_A1 in *T. vaginalis* cells. NlpC_A1-transfected *T. vaginalis* (~10^7^ cells, in total) were pelleted, washed with phosphate-saline buffer (PBS) and fixed with cold methanol for 10 min. Methanol-fixed cells were washed three times with PBS and incubated with 3% bovine serum albumin (BSA) in PBS for 30 min. Cells were pelleted and resuspended with the same 3% BSA in PBS but containing the primary anti-HA (Covance) at 1:1,000 dilution. After 1 hour of incubation, cells were washed three times with 3% BSA in PBS and resuspended in this solution containing Alexa Fluor-conjugated secondary antibody (Invitrogen) at 1:5,000 dilution. After 1 hour of incubation and additional three washes, ~ 10 μL of cells were spotted on a microscope slide along with ~20 ul of ProLong Gold antifade reagent containing 4’-6-diamidino-2-phenylindole or DAPI (Invitrogen) and covered with a coverslip. Images were taken using a Nikon Ni-U microscope equipped with a Spot Pursuit Slider camera (greyscale, cooled, 1.4 megapixel, with colour by filter) and analyzed by its built-in Spot software. Non-transfected *T. vaginalis* cells, used as negative control, did not produce any detectable signal above the background with the anti-HA antibody.

Alternatively, a cell fractionation protocol [89] was applied to NlpC_A1-transfected *T. vaginalis*. A total of ~10^8^ cells were pelleted and washed firstly with PBS followed by buffer A (10 mM HEPES, 1.5 mM MgCl2, 10 mM KCL, 0.5 mM DTT), 1 mL each. The cell pellet was then resuspended in 1 mL of buffer A per gram of cells and incubated for 10 min. Cells were burst by 5-10 passages through a 25G-needle syringe. The cell lysate was taken through steps of differential-speed centrifugation [89]. Protein fractions corresponding to organelles (including nuclei), cytosol and cell membrane were obtained. All procedures were carried out on ice-temperature with all solutions containing protease inhibitor cocktail (complete mini, Roche). In addition to these crude cell fraction protein extracts, a whole-cell protein extract was prepared in parallel. Importantly, each fraction was brought up to equivalent volumes as to the whole cell extract to allow comparisons after SDS-PAGE and Western blot analyses. Replicas of SDS-PAGE gels were stained with Coomassie blue and blotted to PVDF membranes. Western blots were probed with either a mouse monoclonal anti-HA (Covance) or a rabbit polyclonal anti-ferredoxin (kindly donated by Patricia Johnson, UCLA), an organellar marker for *T. vaginalis* which recognizes a typical hydrogenosomal protein. Mouse- and rabbit-specific secondary antibodies conjugated to horseradish peroxidase were used for chemiluminescent detection, as recommended (ThermoFisher Scientific), with the imager Fuji LAS-4000.

## Acknowledgements

We are grateful for the financial support received from the different funders: the Health Research Council of New Zealand, the Maurice & Phyllis Paykel Trust, the Faculty of Science Research Development Fund (University of New Zealand), the Royal Society of New Zealand and the Wellcome Trust. AS-B thanks Patricia Johnson (UCLA) for kindly donating the empty MasterNeo plasmid and the anti-ferredoxin antibody. AS-B thanks Patricia Johnson and her research group members for invaluable constructive criticisms and inputs on the detailed aspects of this research. The authors want to acknowledge the helpful assistance from technical team of the DNA sequencing core facility at the University of Auckland. Finally, the authors also want to acknowledge the NZ synchrotron group for access to SAXS and MX beamlines at the Australian Synchrotron.

---

**S1 Fig.** Sequence comparison of *T. vaginalis* NlpC/P60 proteins. (A) Multiple sequence alignment of *T. vaginalis* NplC/P60 proteins with selected homologues. The alignment is focused on the conserved NlpC/P60 domain. The positions of residues for the protein encoded by TVAG_119910 are indicated at the beginning of each block of the alignment. The residues of the catalytic Cys-His-His triad are boxed showing complete conservation among aligned sequences. The position of indels (dashes) for the four *T. vaginalis* sequences from clan A are distinct from the five sequences from clan B and are also differentially shared with the sequences from the phage infecting *Clostridium difficile* (accession YP_009217650.1) and three bacteria (*Drancourtella* sp. accession WP_087169682.1; *Streptosporangium canum*, accession SFL23765.1 and *Clostridium bifermentans*, accession EQK42778.1) that are also clustering in the phylogenetic tree with respectively *T. vaginalis* sequences from clan A and clan B (see Fig 1). This further supports two distinct LGT events into the *Trichomonas* lineage for these NlpC/P60 encoding genes, which were followed by gene duplications events leading to a total of nine TvNlpC/P60 genes. (B) Comparison of the structural organisation of the nine *T. vaginalis* NlpC/P60 and the other six eukaryotic NlpC/P60 proteins included in the phylogenetic analyses (Fig 1C, S1 table).

**S2 Fig.** Small angle X-ray scattering analysis of NlpC_A1 and NlpC_A2. Scattering data for **(A)** NlpC_A1 and **(B)** NlpC_A2. A dilution series was collected for each protein and shows no concentration dependence in the scattering profile. Guinier analysis for each concentration is shown inset. The pair-distribution function p(r) for **(C)** NlpC_A1 and **(D)** NlpC_A2. Comparison of the observed scattering profile with the theoritical scattering from the structure of **(E)** NlpC_A1 and **(F)** NlpC_A2.

**S3 Fig.** Mutant NlpC proteins are inactive against *E. coli* PG. HPLC chromatograms from peptidoglycan cleavage assays with NlpC mutant proteins (10 μM) and control (no enzyme) are indicated. Peaks were assigned by comparison with published literature. No detectable activity was observed for any of the mutant proteins.

**S4 Fig.** Cell fractionation indicates that the HA-tagged NlpC_A1 is on the membrane of *T. vaginalis*. Cells transfected with a plasmid expressing the HA-tagged NlpC_A1 were incubated in a hypo-osmostic buffer and broken up by mechanic lysis and cellular fractions were obtained by differential centrifugation (see Methods). Cell fractions were obtained as T (total, i.e. whole cell with no fractionation), O (organelles, including nuclei), C (cytosol) and M (membrane). Samples were loaded on SDS-PAGE gels at equivalent volumes, along with a molecular weight marker (MW), for coomassie staining (first panel) and Western blots. Western blots were probed with the primary antibodies anti-HA (second panel) and anti-ferredoxin (third panel), as indicated. The primary anti-ferredoxin (anti-Fd) detects the hydrogenosomal ferredoxin (a clear indicator of an organellar fraction for *T. vaginalis* cell fractionation) and it was kindly provided by Patricia Johnson (UCLA). Secondary antibodies conjugated with horseradish peroxidase were used for chemiluminescent detection. HA-tagged NlpC_A1 is found in fraction M, indicating membrane localization. This fraction is clean from unbroken cells since no signal with anti-ferredoxin antibody can be detected. However, fraction O still carries signal for HA-tagged NlpC_A1 which may come from intact cells carried over to this fraction due to partial cell lysis.

**S5 Fig.** *T. vaginalis* virtually eliminates *E. coli* from mixed cultures when over-expressing NlpC_A1. *E. coli* DH5α was incubated with NlpC_A1-transfected *T. vaginalis* or with non-transfected *T. vaginalis* G3, as indicated, at a cell ratio of 1:10 (bacteria:protozoan) for 1, 2, 4 and 6 hours. While *E. coli* thrived with non-transfected G3 throughout the course of the experiment, these bacteria were virtually eliminated with transfected *T. vaginalis* over-expressing NlpC_A1 in the first hour or co-incubation.

**S6 Fig.** GC% plot for the nine scaffolds encoding the TvNlpC/P60 proteins. For each scaffold the graph of (i) the negative z’-score and the (ii) GC% composition of each segments are shown. Blue dots correspond to the start and stop codons for the ORF encoding the TvNlpC/P60 protein. The green square(s) indicate the position of the inferred segmentation points between major GC% transitions when present. All GC% values for identified segments are shown on each graph and these are also listed in S1 Table. The individual scaffolds were analysed with the following settings following the instructions provided at the GC-profile server: <u>Settings 1:</u> GC_plots_TVAG_042760_DS113937_2 --- 43,359 bp; GC_plots_TVAG_411960_DS113699_2 --- 63,570 bp <u>with</u> Halting parameter = 10.00 <u>and</u> Minimum length = 100 bp. <u>Settings 2:</u> GC_plots_TVAG_252970_DS113529_2 --- 84,867 bp; GC_plots_TVAG_324990_DS113569_2 --- 79,056 bp <u>with</u> Halting parameter = 50.00 <u>and</u> Minimum length = 100 bp<u>. Settings 3:</u> GC_plots_TVAG_051010_DS113369_2 --- 122,255 bp; GC_plots_TVAG_119910_DS113177_2 --- 584,929 bp; GC_plots_TVAG_209010_DS113253_2 --- 190,068 bp; GC_plots_TVAG_393610_DS113268_2 --- 177,036 bp; GC_plots_TVAG_457240_DS113186_2 --- 405,446 bp<u>; with</u> Halting parameter = 50.00 and Minimum length = 1000 bp.

**S7 Fig.** The bacteriolytic effect of the purified recombinant NlpC_A1 enzyme and lysozyme on *E. coli* DH5α. Bacteria (5 × 10^5^ cell/ml) were incubated with increasing concentrations of the purified recombinant NlpC_A1 wild-type (blue line) and C179S mutant (orange line) in [10 mM Tris-HCl pH 7.5, 100 mM NaCl and 1 mM EDTA] at 37 °C for 1 hour. As a positive control, bacteria were incubated with lysozyme (black line) at the same concentrations: 12.5, 25, 50 and 100 μM. As a negative control, bacteria were incubated with buffer only (no enzyme). Following incubation, bacteria were plated on LB-agar with appropriate dilution for cfu counting. Reduction on cfu was calculated as a percentage in comparison to the mock treatment (negative control). While lysozyme promotes cfu reduction due to its known bactericidal activity, purified NlpC_A1 even at the highest concentration does not have any effect. Therefore, *T. vaginalis* NlpC_A1 is not bacteriolytic.

**S8 Fig.** Treefile used for the Approximate Unbiased (AU) topological test. The text file lists the eight trees (in NEWICK format) used for the AU topological test. The results of the AU test and tree descriptions are listed in S5 Table.

**S1 Table.** Overview of *T. vaginalis* NlpC/P60 genes, gene expression data and protein features. **Tv**NlpC/P60 genes/proteins were sorted by clusters A (in blue) and B (in green).

**S2 Table.** Taxonomic BlastP report for query NlpC_A1/TVAG_119910/XP_001276902 against the NCBI nr protein database.

**S3 Table.** Taxonomic BlastP report for query NlpC_B1/TVAG_393610/XP_001326856 against the NCBI nr protein database.

**S4 Table.** Taxonomic DELTA-Blast report for query NlpC_A1/TVAG_119910/XP_001276902 against the NCBI nr protein database restricted to eukaryotes.

**S5 Table.** Approximate Unbiased (AU) tree topological test for hypotheses of TvNlpC/P60 phylogenetic relationships. Tree topologies derived from unconstrained analyses using different amino acid evolutionary models were tested against each other and against tree topologies derived from four constrained analyses testing specific relationships for the TvNlpC/P60 entries, as described in the main text.

**S6 Table.** Data collection and refinement statistics. Values in parenthesis represent the highest resolution shell.

**S7 Table.** Small-angle X-ray scattering data analysis.

## References

1. WHO. Global health sector strategy on sexually transmitted infections 2016-2021 [Internet]. Who. World Health Organization; 2016. Available: http://www.who.int/reproductivehealth/publications/rtis/ghss-stis/en/

2. Swygard H, Seña AC, Hobbs MM, Cohen MS. Trichomoniasis: clinical manifestations, diagnosis and management. Sex Transm Infect. 2004;80: 91–95. doi:10.1136/sti.2003.005124

3. Cotch MF, Pastorek JG, Nugent RP, Hillier SL, Gibbs RS, Martin DH, et al. Trichomonas vaginalis associated with low birth weight and preterm delivery. Sex Transm Dis. 1997;24: 353–360. doi:10.1097/00007435-199707000-00008

4. Hardy P, Nell EE, Spence M, Hardy J, Graham D, Rosenbaum R. Prevalence of Six Sexually Transmitted Disease Agents Among Pregnant Inner-City Adolescents and Pregnancy Outcome. Lancet. 1984;324: 333–337. doi:10.1016/S0140-6736(84)92698-9

5. Minkoff H, Grunebaum AN, Schwarz RH, Feldman J, Cummings M, Crombleholme W, et al. Risk factors for prematurity and premature rupture of membranes: A prospective study of the vaginal flora in pregnancy. Am J Obstet Gynecol. Mosby; 1984;150: 965–972. doi:10.1016/0002-9378(84)90392-2

6. Gram IT, Macaluso M, Churchill J, Stalsberg H. Trichomonas vaginalis (TV) and human papillomavirus (HPV) infection and the incidence of cervical intraepithelial neoplasia (CIN) grade III. Cancer Causes Control. 1992;3: 231–6. doi:10.1007/BF00124256.

7. Stark JR, Judson G, Alderete JF, Mundodi V, Kucknoor AS, Giovannucci EL, et al. Prospective study of trichomonas vaginalis infection and prostate cancer incidence and mortality: Physicians’ health study. J Natl Cancer Inst. 2009;101: 1406–1411. doi:10.1093/jnci/djp306

8. Laga M, Manoka A, Kivuvu M, Malele B, Tuliza M, Nzila N, et al. Non-ulcerative sexually transmitted diseases as risk factors for HIV-1 transmission in women: results from a cohort study. AIDS. 1993;7: 95–102.

9. Fichorova RN, Buck OR, Yamamoto HS, Fashemi T, Dawood HY, Fashemi B, et al. The villain team-up or how Trichomonas vaginalis and bacterial vaginosis alter innate immunity in concert. Sex Transm Infect. 2013;89: 460–466. doi:10.1136/sextrans-2013-051052

10. Phukan N, Parsamand T, Brooks AES, Nguyen TNM, Simoes-Barbosa A. The adherence of Trichomonas vaginalis to host ectocervical cells is influenced by lactobacilli. Sex Transm Infect. 2013;89: 455–459. doi:10.1136/sextrans-2013-051039

11. Bär A-K, Phukan N, Pinheiro J, Simoes-Barbosa A. The Interplay of Host Microbiota and Parasitic Protozoans at Mucosal Interfaces: Implications for the Outcomes of Infections and Diseases. PLoS Negl Trop Dis. 2015;9: e0004176. doi:10.1371/journal.pntd.0004176

12. Brotman RM, Bradford LL, Conrad M, Gajer P, Aul K, Peralta L, et al. Association between Trichomonas vaginalis and vaginal bacterial community composition among reproductive-age women. Sex Transm Dis. 2012;39: 807–812. doi:10.1097/OLQ.0b013e3182631c79

13. Maritz JM, Land KM, Carlton JM, Hirt RP. What is the importance of zoonotic trichomonads for human health? Trends Parasitol. 2014;30: 333–341. doi:10.1016/j.pt.2014.05.005

14. Carlton JM, Hirt RP, Silva JC, Delcher AL, Schatz M, Zhao Q, et al. Draft Genome Sequence of the Sexually Transmitted Pathogen Trichomonas vaginalis. Science (80-). 2007;315: 207–212. doi:10.1126/science.1132894

15. Hirt RP, Alsmark C, Embley TM. Lateral gene transfers and the origins of the eukaryote proteome: a view from microbial parasites. Curr Opin Microbiol. 2015;23: 155–162. doi:10.1016/j.mib.2014.11.018

16. Huang K, Chen Y-YM, Fang Y, Cheng W-H, Cheng C, Chen Y-C, et al. Adaptive responses to glucose restriction enhance cell survival, antioxidant capability, and autophagy of the protozoan parasite Trichomonas vaginalis. Biochim Biophys Acta - Gen Subj. 2014;1840: 53–64. doi:10.1016/j.bbagen.2013.08.008

17. Gould SB, Woehle C, Kusdian G, Landan G, Tachezy J, Zimorski V, et al. Deep sequencing of Trichomonas vaginalis during the early infection of vaginal epithelial cells and amoeboid transition. Int J Parasitol. Australian Society for Parasitology Inc.; 2013;43: 707–719. doi:10.1016/j.ijpara.2013.04.002

18. Alsmark C, Foster PG, Sicheritz-Ponten T, Nakjang S, Martin Embley T, Hirt RP. Patterns of prokaryotic lateral gene transfers affecting parasitic microbial eukaryotes. Genome Biol. 2013;14: R19. doi:10.1186/gb-2013-14-2-r19

19. Nikoh N, McCutcheon JP, Kudo T, Miyagishima SY, Moran NA, Nakabachi A. Bacterial genes in the aphid genome: Absence of functional gene transfer from Buchnera to its host. PLoS Genet. 2010;6: e1000827. doi:10.1371/journal.pgen.1000827

20. Schönknecht G, Weber APM, Lercher MJ. Horizontal gene acquisitions by eukaryotes as drivers of adaptive evolution. BioEssays. 2014;36: 9–20. doi:10.1002/bies.201300095

21. Soucy SM, Huang J, Gogarten JP. Horizontal gene transfer: building the web of life. Nat Rev Genet. 2015;16: 472–482. doi:10.1038/nrg3962

22. Husnik F, McCutcheon JP. Functional horizontal gene transfer from bacteria to eukaryotes. Nat Rev Microbiol. 2017;16: 67–79. doi:10.1038/nrmicro.2017.137

23. Gladyshev EA, Meselson M, Arkhipova IR. Massive Horizontal Gene Transfer in Bdelloid Rotifers. Science (80-). 2008;320: 1210–1213. doi:10.1126/science.1156407

24. Nikoh N, Nakabachi A. Aphids acquired symbiotic genes via lateral gene transfer. BMC Biol. 2009;7: 12. doi:10.1186/1741-7007-7-12

25. Metcalf JA, Funkhouser-Jones LJ, Brileya K, Reysenbach A, Bordenstein SR. Antibacterial gene transfer across the tree of life. Elife. 2014;3: 1–18. doi:10.7554/eLife.04266

26. Chou S, Daugherty MD, Peterson SB, Biboy J, Yang Y, Jutras BL, et al. Transferred interbacterial antagonism genes augment eukaryotic innate immune function. Nature. 2015;518: 98–101. doi:10.1038/nature13965

27. Ohnishi R, Ishikawa S, Sekiguchi J. Peptidoglycan hydrolase LytF plays a role in cell separation with CwlF during vegetative growth of Bacillus subtilis. J Bacteriol. 1999;181: 3178–3184. Available: http://www.ncbi.nlm.nih.gov/pubmed/10322020

28. Anantharaman V, Aravind L. Evolutionary history, structural features and biochemical diversity of the NlpC/P60 superfamily of enzymes. Genome Biol. 2003;4: R11. doi:10.1186/gb-2003-4-2-r11

29. Aurrecoechea C, Brestelli J, Brunk BP, Carlton JM, Dommer J, Fischer S, et al. GiardiaDB and TrichDB: integrated genomic resources for the eukaryotic protist pathogens Giardia lamblia and Trichomonas vaginalis. Nucleic Acids Res. 2009;37: D526–D530. doi:10.1093/nar/gkn631

30. Dyrløv Bendtsen J, Nielsen H, von Heijne G, Brunak S. Improved Prediction of Signal Peptides: SignalP 3.0. J Mol Biol. 2004;340: 783–795. doi:10.1016/J.JMB.2004.05.028

31. Rancurel C, Legrand L, Danchin EGJ. Alienness: Rapid detection of candidate horizontal gene transfers across the tree of life. Genes (Basel). 2017;8. doi:10.3390/genes8100248

32. Benchimol M, de Almeida LGP, Vasconcelos AT, de Andrade Rosa I, Reis Bogo M, Kist LW, et al. Draft Genome Sequence of Tritrichomonas foetus Strain K. Genome Announc. 2017;5: e00195–17. doi:10.1128/genomeA.00195-17

33. Brown CT, Hug LA, Thomas BC, Sharon I, Castelle CJ, Singh A, et al. Unusual biology across a group comprising more than 15% of domain Bacteria. Nature. 2015;523: 208–211. doi:10.1038/nature14486

34. Clarke M, Lohan AJ, Liu B, Lagkouvardos I, Roy S, Zafar N, et al. Genome of *Acanthamoeba castellanii* highlights extensive lateral gene transfer and early evolution of tyrosine kinase signaling. Genome Biol. 2013;14: R11. doi:10.1186/gb-2013-14-2-r11

35. Tang VH, Stewart GA, Chang BJ. *Dermatophagoides pteronyssinus lytFM* encoding an NlpC/P60 endopeptidase is also present in mite-associated bacteria that express LytFM variants. FEBS Open Bio. 2017;7: 1267–1280. doi:10.1002/2211-5463.12263

36. Strese Å, Backlund A, Alsmark C. A recently transferred cluster of bacterial genes in Trichomonas vaginalis - lateral gene transfer and the fate of acquired genes. BMC Evol Biol. 2014;14: 119. doi:10.1186/1471-2148-14-119

37. Drenth J, Jansonius JN, Koekoek R, Swen HM, Wolthers BG. Structure of Papain. Nature. 1968;218: 929–932. doi:10.1038/218929a0

38. Typas A, Banzhaf M, Gross CA, Vollmer W. From the regulation of peptidoglycan synthesis to bacterial growth and morphology. Nat Rev Microbiol. 2011;10: 123–136. doi:10.1038/nrmicro2677

39. Baker NA, Sept D, Joseph S, Holst MJ, McCammon JA. Electrostatics of nanosystems: Application to microtubules and the ribosome. Proc Natl Acad Sci. 2001;98: 10037–10041. doi:10.1073/pnas.181342398

40. Wong JEMM, Midtgaard SR, Gysel K, Thygesen MB, Sørensen KK, Jensen KJ, et al. An intermolecular binding mechanism involving multiple LysM domains mediates carbohydrate recognition by an endopeptidase. Acta Crystallogr Sect D Biol Crystallogr. 2015;71: 592–605. doi:10.1107/S139900471402793X

41. Krissinel E, Henrick K. Secondary-structure matching (SSM), a new tool for fast protein structure alignment in three dimensions. Acta Crystallogr Sect D Biol Crystallogr. 2004;60: 2256–2268. doi:10.1107/S0907444904026460

42. Xu Q, Abdubek P, Astakhova T, Axelrod HL, Bakolitsa C, Cai X, et al. Structure of the γ-D -glutamyl- L -diamino acid endopeptidase YkfC from *Bacillus cereus* in complex with L -Al. Acta Crystallogr Sect F Struct Biol Cryst Commun. 2010;66: 1354–1364. doi:10.1107/S1744309110021214

43. Casadaban MJ, Cohen SN. Analysis of gene control signals by DNA fusion and cloning in Escherichia coli. J Mol Biol. 1980;138: 179–207. doi:10.1016/0022-2836(80)90283-1

44. Meberg BM, Sailer FC, Nelson DE, Young KD. Reconstruction of Escherichia coli mrcA (PBP 1a) Mutants Lacking Multiple Combinations of Penicillin Binding Proteins. J Bacteriol. 2001;183: 6148–6149. doi:10.1128/JB.183.20.6148-6149.2001

45. Lawrence JG, Ochman H. Amelioration of Bacterial Genomes: Rates of Change and Exchange. J Mol Evol. 1997;44: 383–397. doi:10.1007/PL00006158

46. Bertsche U, Breukink E, Kast T, Vollmer W. In Vitro Murein (Peptidoglycan) Synthesis by Dimers of the Bifunctional Transglycosylase-Transpeptidase PBP1B from Escherichia coli. J Biol Chem. 2005;280: 38096–38101. doi:10.1074/jbc.M508646200

47. Vollmer W, Joris B, Charlier P, Foster S. Bacterial peptidoglycan (murein) hydrolases. FEMS Microbiol Rev. 2008;32: 259–286. doi:10.1111/j.1574-6976.2007.00099.x

48. de Miguel N, Lustig G, Twu O, Chattopadhyay A, Wohlschlegel JA, Johnson PJ. Proteome Analysis of the Surface of Trichomonas vaginalis Reveals Novel Proteins and Strain-dependent Differential Expression. Mol Cell Proteomics. 2010;9: 1554–1566. doi:10.1074/mcp.M000022-MCP201

49. Huang K-Y, Huang P-J, Ku F-M, Lin R, Alderete JF, Tang P. Comparative Transcriptomic and Proteomic Analyses of Trichomonas vaginalis following Adherence to Fibronectin. Infect Immun. 2012;80: 3900–3911. doi:10.1128/IAI.00611-12

50. Twu O, de Miguel N, Lustig G, Stevens GC, Vashisht AA, Wohlschlegel JA, et al. Trichomonas vaginalis Exosomes Deliver Cargo to Host Cells and Mediate Host:Parasite Interactions. Petri WA, editor. PLoS Pathog. 2013;9: e1003482. doi:10.1371/journal.ppat.1003482

51. Ragland SA, Criss AK. From bacterial killing to immune modulation: Recent insights into the functions of lysozyme. Bliska JB, editor. PLOS Pathog. 2017;13: e1006512. doi:10.1371/journal.ppat.1006512

52. Bard E, Laibe S, Bettinger D, Riethmuller D, Biichlé S, Seilles E, et al. New Sensitive Method for the Measurement of Lysozyme and Lactoferrin for the Assessment of Innate Mucosal Immunity. Part I: Time-Resolved Immunofluorometric Assay in Serum and Mucosal Secretions. Clin Chem Lab Med. 2003;41: 127–133. doi:10.1515/CCLM.2003.021

53. Chou S, Bui NK, Russell AB, Lexa KW, Gardiner TE, LeRoux M, et al. Structure of a Peptidoglycan Amidase Effector Targeted to Gram-Negative Bacteria by the Type VI Secretion System. Cell Rep. 2012;1: 656–664. doi:10.1016/j.celrep.2012.05.016

54. Pazos M, Peters K, Vollmer W. Robust peptidoglycan growth by dynamic and variable multi-protein complexes. Curr Opin Microbiol. 2017;36: 55–61. doi:10.1016/j.mib.2017.01.006

55. Wolf AJ, Underhill DM. Peptidoglycan recognition by the innate immune system. Nat Rev Immunol. 2018;18: 243–254. doi:10.1038/nri.2017.136

56. Chu H, Mazmanian SK. Innate immune recognition of the microbiota promotes host-microbial symbiosis. Nat Immunol. 2013;14: 668–75. doi:10.1038/ni.2635

57. Fichorova R, Fraga J, Rappelli P, Fiori PL. Trichomonas vaginalis infection in symbiosis with Trichomonasvirus and Mycoplasma. Res Microbiol. 2017;168: 882–891. doi:10.1016/J.RESMIC.2017.03.005

58. Finn RD, Attwood TK, Babbitt PC, Bateman A, Bork P, Bridge AJ, et al. InterPro in 2017-beyond protein family and domain annotations. Nucleic Acids Res. 2017;45: D190–D199. doi:10.1093/nar/gkw1107

59. Käll L, Krogh A, Sonnhammer ELL. Advantages of combined transmembrane topology and signal peptide prediction-the Phobius web server. Nucleic Acids Res. 2007;35: W429–W432. doi:10.1093/nar/gkm256

60. Nielsen H. Predicting Secretory Proteins with SignalP. Protein Function Prediction Methods in Molecular Biology. 2017. pp. 59–73. doi:10.1007/978-1-4939-7015-5_6

61. Viklund H, Bernsel A, Skwark M, Elofsson A. SPOCTOPUS: A combined predictor of signal peptides and membrane protein topology. Bioinformatics. 2008;24: 2928–2929. doi:10.1093/bioinformatics/btn550

62. Capella-Gutierrez S, Silla-Martinez JM, Gabaldon T. trimAl: a tool for automated alignment trimming in large-scale phylogenetic analyses. Bioinformatics. 2009;25: 1972–1973. doi:10.1093/bioinformatics/btp348

63. Lefort V, Longueville J, Gascuel O. SMS: Smart Model Selection in PhyML. Mol Biol Evol. 2017;34: 2422–2424. doi:10.1093/molbev/msx149

64. Le SQ, Dang CC, Gascuel O. Modeling protein evolution with several amino acid replacement matrices depending on site rates. Mol Biol Evol. 2012;29: 2921–2936. doi:10.1093/molbev/mss112

65. Quang LS, Gascuel O, Lartillot N. Empirical profile mixture models for phylogenetic reconstruction. Bioinformatics. 2008;24: 2317–2323. doi:10.1093/bioinformatics/btn445

66. Guindon S, Dufayard J-F, Lefort V, Anisimova M, Hordijk W, Gascuel O. New Algorithms and Methods to Estimate Maximum-Likelihood Phylogenies: Assessing the Performance of PhyML 3.0. Syst Biol. 2010;59: 307–321. doi:10.1093/sysbio/syq010

67. Gouy M, Guindon S, Gascuel O. SeaView Version 4: A Multiplatform Graphical User Interface for Sequence Alignment and Phylogenetic Tree Building. Mol Biol Evol. 2010;27: 221–224. doi:10.1093/molbev/msp259

68. Nguyen LT, Schmidt HA, Von Haeseler A, Minh BQ. IQ-TREE: A fast and effective stochastic algorithm for estimating maximum-likelihood phylogenies. Mol Biol Evol. 2015;32: 268–274. doi:10.1093/molbev/msu300

69. Trifinopoulos J, Nguyen LT, von Haeseler A, Minh BQ. W-IQ-TREE: a fast online phylogenetic tool for maximum likelihood analysis. Nucleic Acids Res. 2016;44: W232–W235. doi:10.1093/nar/gkw256

70. Shimodaira H. An approximately unbiased test of phylogenetic tree selection. Syst Biol. 2002;51: 492–508. doi:10.1080/10635150290069913

71. Letunic I, Bork P. Interactive tree of life (iTOL) v3: an online tool for the display and annotation of phylogenetic and other trees. Nucleic Acids Res. 2016;44: W242–W245. doi:10.1093/nar/gkw290

72. Gao F, Zhang CT. GC-Profile: A web-based tool for visualizing and analyzing the variation of GC content in genomic sequences. Nucleic Acids Res. 2006;34: 686–691. doi:10.1093/nar/gkl040

73. Puigbò P, Bravo IG, Garcia-Vallve S. CAIcal: A combined set of tools to assess codon usage adaptation. Biol Direct. 2008;3: 1–8. doi:10.1186/1745-6150-3-38

74. Clark CG, Diamond LS. Methods for Cultivation of Luminal Parasitic Protists of Clinical Importance. Clin Microbiol Rev. 2002;15: 329–341. doi:10.1128/CMR.15.3.329-341.2002

75. Brooks AES, Parsamand T, Kelly RW, Simoes-Barbosa A. An improved quantitative method to assess adhesive properties of Trichomonas vaginalis to host vaginal ectocervical cells using flow cytometry. J Microbiol Methods. 2013;92: 73–78. doi:10.1016/j.mimet.2012.10.011

76. Pfaffl MW. Relative expression software tool (REST(C)) for group-wise comparison and statistical analysis of relative expression results in real-time PCR. Nucleic Acids Res. 2002;30: 36e–36. doi:10.1093/nar/30.9.e36

77. Delgadillo MG, Liston DR, Niazi K, Johnson PJ. Transient and selectable transformation of the parasitic protist Trichomonas vaginalis. Proc Natl Acad Sci. 1997;94: 4716–4720. doi:10.1073/pnas.94.9.4716

78. Aslanidis C, de Jong PJ. Ligation-independent cloning of PCR products (LIC-PCR). Nucleic Acids Res. 1990;18: 6069–6074. doi:10.1093/nar/18.20.6069

79. Studier FW. Protein production by auto-induction in high-density shaking cultures. Protein Expr Purif. 2005;41: 207–234. doi:10.1016/j.pep.2005.01.016

80. Gorrec F. The MORPHEUS protein crystallization screen. J Appl Crystallogr. International Union of Crystallography; 2009;42: 1035–1042. doi:10.1107/S0021889809042022

81. Moreland N, Ashton R, Baker HM, Ivanovic I, Patterson S, Arcus VL, et al. A flexible and economical medium-throughput strategy for protein production and crystallization. Acta Crystallogr Sect D Biol Crystallogr. 2005;61: 1378–1385. doi:10.1107/S0907444905023590

82. Sheldrick GM. Experimental phasing with SHELXC / D / E : combining chain tracing with density modification. Acta Crystallogr Sect D Biol Crystallogr. International Union of Crystallography; 2010;66: 479–485. doi:10.1107/S0907444909038360

83. Potterton E, Briggs P, Turkenburg M, Dodson E. A graphical user interface to the CCP4 program suite. Acta Crystallogr Sect D Biol Crystallogr. 2003;59: 1131–1137. doi:10.1107/S0907444903008126

84. Adams PD, Baker D, Brunger AT, Das R, DiMaio F, Read RJ, et al. Advances, Interactions, and Future Developments in the CNS, Phenix, and Rosetta Structural Biology Software Systems. Annu Rev Biophys. 2013;42: 265–287. doi:10.1146/annurev-biophys-083012-130253

85. Emsley P, Lohkamp B, Scott WG, Cowtan K. Features and development of Coot. Acta Crystallogr Sect D Biol Crystallogr. 2010;66: 486–501. doi:10.1107/S0907444910007493

86. McCoy AJ. Solving structures of protein complexes by molecular replacement with Phaser. Acta Crystallographica Section D: Biological Crystallography. 2006. pp. 32–41. doi:10.1107/S0907444906045975

87. Bui NK, Gray J, Schwarz H, Schumann P, Blanot D, Vollmer W. The peptidoglycan sacculus of Myxococcus xanthus has unusual structural features and is degraded during glycerol-induced myxospore development. J Bacteriol. 2009;191: 494–505. doi:10.1128/JB.00608-08

88. Russell AB, Singh P, Brittnacher M, Bui NK, Hood RD, Carl MA, et al. A Widespread Bacterial Type VI Secretion Effector Superfamily Identified Using a Heuristic Approach. Cell Host Microbe. 2012;11: 538–549. doi:10.1016/j.chom.2012.04.007

89. Yu Z, Huang Z, Lung M. Subcellular Fractionation of Cultured Human Cell Lines. Bio-Protocol. 2013;3. doi:10.21769/BioProtoc.754

